# Quantifying Mg^2+^ dependence on conformational equilibrium in the two-state 7SK RNA stem-loop 3

**DOI:** 10.64898/2026.04.11.717930

**Authors:** Shilla Owusu Ansah, Momodou B. Camara, Catherine D. Eichhorn

**Affiliations:** Department of Chemistry, University of Nebraska - Lincoln, 639 North 12th St, Lincoln, NE 68588, USA; Nebraska Center for Integrated Biomolecular Communication

**Keywords:** solution NMR spectroscopy, conformational exchange, chemical probing, RNA dynamics, linkage equilibrium, ion-RNA interactions, isothermal titration calorimetry

## Abstract

RNA structural heterogeneity is increasingly recognized as essential for RNA function, yet quantitative understanding of intrinsic RNA structural dynamics and how cellular conditions modulate these dynamics remains limited. 7SK RNA is an abundant eukaryotic noncoding RNA that assembles with protein cofactors to form the 7SK ribonucleoprotein (RNP), a dynamic complex that regulates transcription elongation. In particular, the 7SK RNA stem-loop 3 (SL3) domain is a critical hub for protein recruitment that is required for 7SK RNP function. We recently discovered that SL3 undergoes exchange between two equally populated yet structurally distinct conformers, named SL3e and SL3a, providing a model system to understand RNA conformational equilibria. Here, we combined quantitative dimethyl sulfate mutational profiling with sequencing (qDMS-MaPseq), solution NMR spectroscopy, and isothermal titration calorimetry (ITC) experiments to gain quantitative insights into SL3 equilibria under varying ionic conditions. We find that Mg^2+^ shifts the SL3 conformational equilibrium from 52%:48% to 67%:33% populations, with ITC experiments showing ∼2-fold higher affinity for the SL3e conformer, providing the thermodynamic basis for this population shift. NMR ^1^H-^1^H NOESY experiments show that Mg^2+^ stabilizes an A-form helical geometry in the distal end of the SL3e conformer that contains multiple noncanonical base pairs. Validating our combined approach, NMR and DMS-MaPseq Mg^2+^-induced perturbation data are highly correlated for nucleobase moieties at the Watson-Crick face (R^2^ = 0.87). From this work, we identify a characteristic DMS reactivity signature for A•C wobble base pairs from DMS-MaPseq MgCl_2_ titration experiments, expanding the utility of chemical probing for detecting noncanonical base pairs. Our findings demonstrate how modest differences in cation affinity between conformational states can modulate RNA population ensembles. More broadly, this work establishes a generalizable thermodynamic framework to quantitatively dissect ion-dependent conformational equilibria in structurally heterogeneous RNAs and advances structural understanding of the essential SL3 regulatory element in the 7SK RNP.

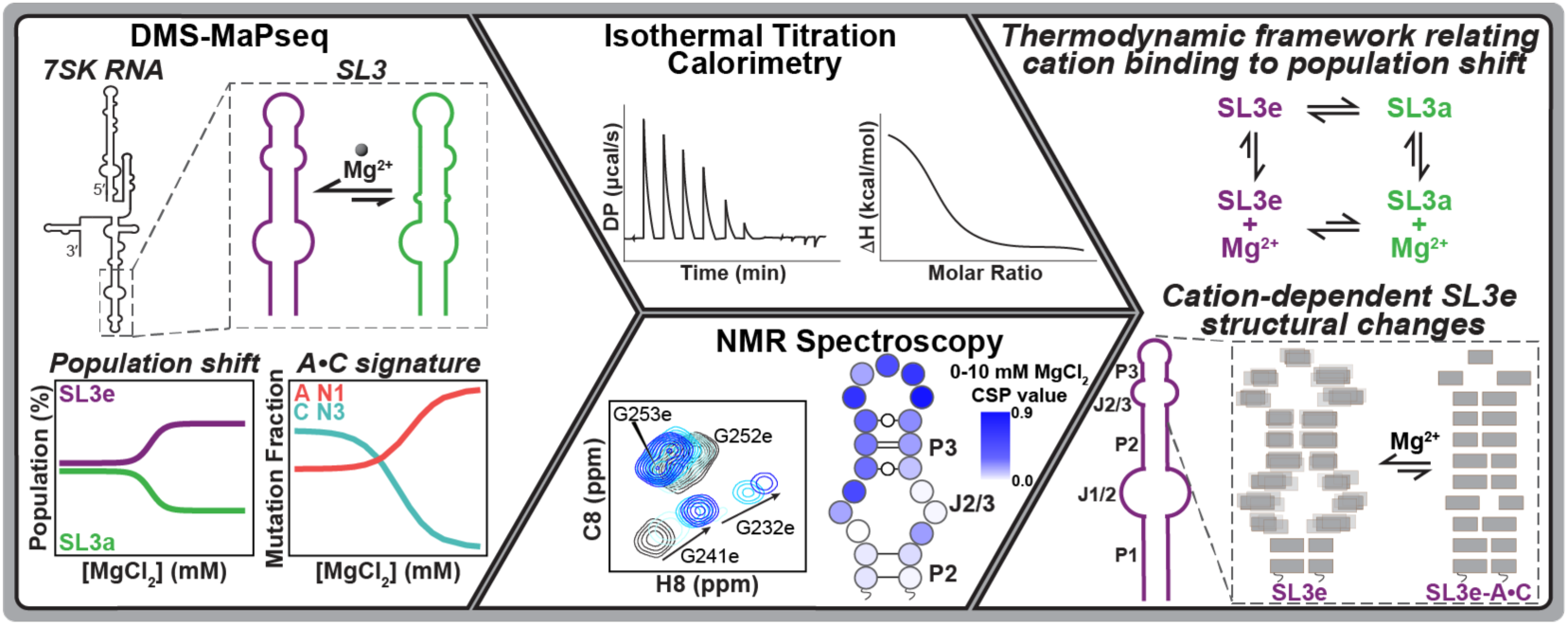

**KEY POINTS:**

- Addition of Mg^2+^ shifts the SL3 two-state conformational equilibrium through preferential association with the SL3e state
- Mg^2+^ association stabilizes an A-form-like geometry in the SL3e upper stem
- DMS reactivity patterns in MgCl_2_ titration experiments provide a signature for identifying A•C wobble base pairs
- DMS-MaPseq mutation fraction perturbation data are correlated with NMR chemical shift perturbation data, validating chemical probing as a structural reporter

## INTRODUCTION

RNA is a structurally plastic biomolecule whose conformational dynamics are essential for diverse cellular functions, including catalysis, gene regulation, and protein recognition (1). Increasing evidence indicates that rather than a single static fold, RNA exists as ensembles of interconverting structures (2–6). Despite growing recognition of the biological significance of RNA structural heterogeneity (3,4,7), quantitative understanding of RNA conformational equilibria and understanding how cellular factors modulate these ensembles remains limited.

At the secondary structure level, RNA is composed of helices, junctions, and apical loops (8). Divalent metal ions play a central role in RNA folding and structural stability (9). Among these, the magnesium cation (Mg^2+^) is particularly important due to its high cellular abundance (∼1 mM free concentration) (10) and strong charge density (11). Mg^2+^ stabilizes RNA structures through diffuse electrostatic and water-mediated interactions and/or site-specific coordination to electronegative pockets within structured motifs (12,13). Cation binding frequently occurs at electronegative pockets created by noncanonical motifs, including G•U wobble base-pairs, internal loops, and tertiary contacts (14–16). By neutralizing backbone charge and stabilizing local interactions, Mg^2+^ promotes formation of compact RNA folds and tertiary structures (17). Previous studies of the P5abc subdomain of the Tetrahymena group I intron have shown that Mg^2+^ modulates populations of RNA conformational states in the P5c hairpin to induce tertiary folding (18,19). While thermodynamic frameworks have been developed to describe Mg^2+^-dependent folding transitions between unfolded and folded RNA states (11), such frameworks are lacking for RNAs ensembles with multiple folded states. This gap limits a quantitative understanding of how Mg^2+^ binding may result in population shifts within the RNA ensemble.

Chemical probing is a rapidly expanding technique to study RNA structure, with applications for protein binding (20), measuring tertiary contacts (21,22), and deconvoluting RNAs with alternate states (23–25). However, there remains a limited understanding to the degree that chemical probing data quantitatively reports on RNA structure, especially for ligand-RNA interactions. NMR is a complementary technique that provides atomic-level information on both RNA structure and conformational dynamics and has been used previously to validate chemical probing data (26,27).

The noncoding 7SK RNA (331-334 nucleotides in humans) is a central regulator of transcription elongation in eukaryotic cells (28,29). 7SK RNA assembles with multiple protein cofactors to form the 7SK ribonucleoprotein (RNP), which controls the activity of the positive transcription elongation factor b (P-TEFb) (30,31). The 7SK RNA folds into four primary domains (32,33), with stem-loop 3 (SL3) serving as a critical hub for recruitment of accessory proteins that lead to release of P-TEFb (34,35). Despite its functional importance, the structural basis underlying SL3 function remains incompletely understood. Using a combined nuclear magnetic resonance (NMR) spectroscopy and dimethyl sulfate mutational profiling with sequencing (DMS-MaPseq) approach, we previously demonstrated that SL3 undergoes conformational exchange between two distinct structural states, termed SL3e and SL3a, that are nearly equally populated at 25 °C (**Fig. 1A**) (27).

**Figure 1.**
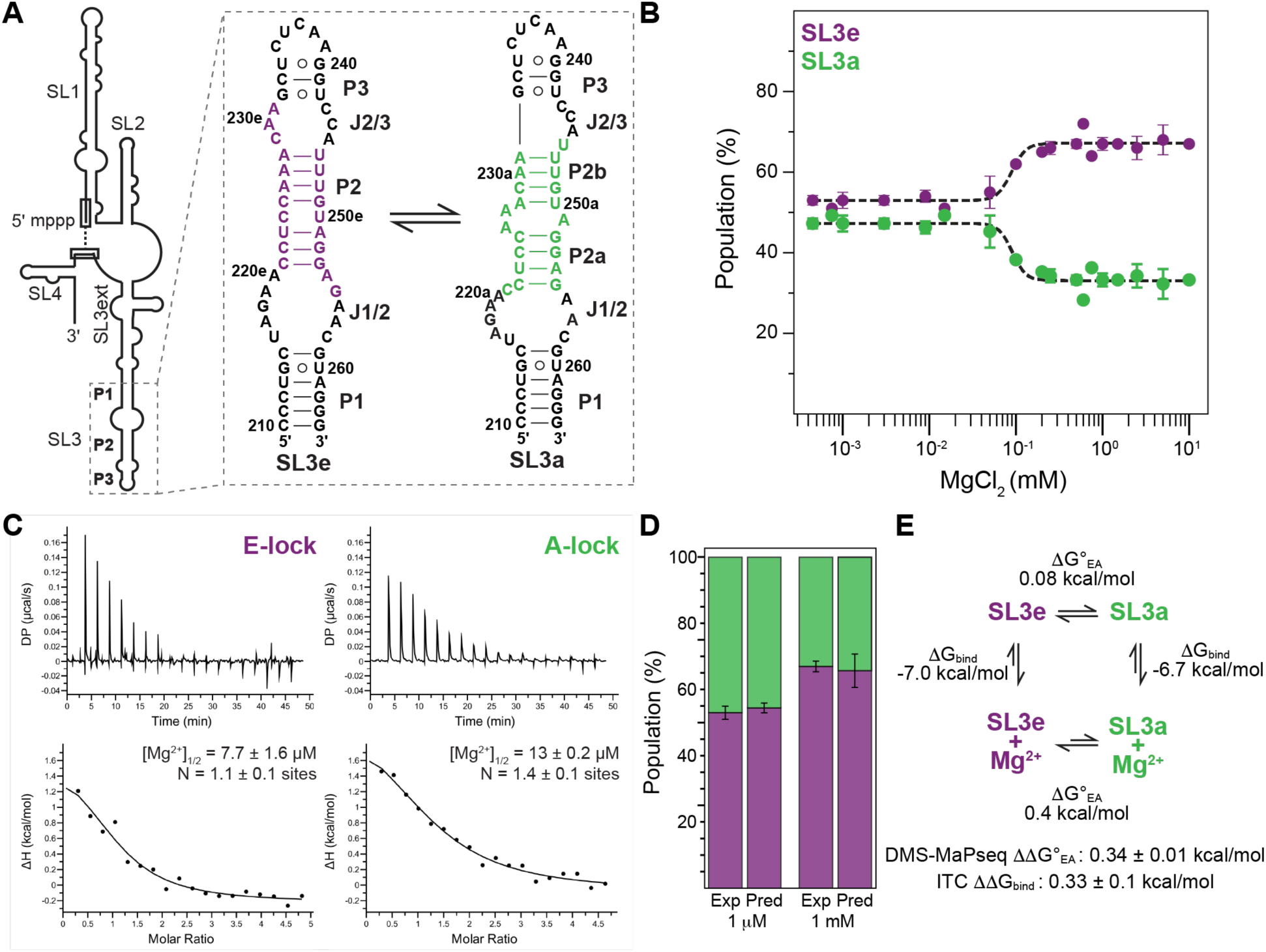
Mg^2+^ shifts the SL3 conformational equilibrium toward the SL3e state through differential cation affinities. **(A)** Secondary structure models of the two conformational states of 7SK RNA SL3. **(B)** Population as a function of MgCl_2_ concentration determined by DREEM clustering of DMS-MaPseq data at 25 °C. Data were fit to a cooperative model yielding a transition midpoint of 90 ± 15 μM Mg^2+^. Error bars represent standard deviation from two independent replicates. **(C)** Isothermal titration calorimetry (ITC) measurements of Mg^2+^ binding to E-lock (left) and A-lock (right) constructs at 25 °C. **(D)** Experimental (Exp) or predicted (Pred) populations of SL3e and SL3a states as a function of MgCl_2_ concentration calculated from a linkage equilibrium model using the ITC-determined binding constants. **(E)** Thermodynamic cycle relating the SL3a-SL3e conformational equilibrium to Mg^2+^ binding, showing that differential cation affinities drive the population shift.

Here we investigate how Mg^2+^ modulates this conformational equilibrium using quantitative DMS-MaPseq (21), solution state NMR spectroscopy, and isothermal titration calorimetry (ITC). We find that physiological Mg^2+^ concentrations shift the equilibrium toward the SL3e conformer through differential association to the upper stem-loop region. Using conformationally-locked constructs, ITC experiments show ∼2-fold higher affinity for the SL3e conformer. In the SL3e conformer, we find that Mg^2+^ stabilizes an A-form-like helical geometry in the distal end of SL3 that is enriched in noncanonical base-pairs. NMR and qDMS-MaPseq Mg^2+^-induced perturbation data are highly correlated for nucleobase moieties at the Watson-Crick face (R^2^ = 0.87). From this work, we identify an A•C wobble base pair signature from qDMS-MaPseq MgCl_2_ titration experiments, expanding the utility of chemical probing for detecting noncanonical base pairs. Our findings demonstrate how modest differences in cation affinity between conformational states can modulate RNA population ensembles. These results provide quantitative insight into how cation association reshapes RNA conformational ensembles and establish an integrated experimental framework for linking ion binding thermodynamics to RNA structural equilibria.

## MATERIALS AND METHODS

### RNA sample preparation

Chemically synthesized DNA templates of SL3 domain constructs were purchased from Integrated DNA Technologies (IDT) (**Supplementary Table S1-2**). RNA samples were prepared by *in vitro* transcription using T7 RNA polymerase (Addgene #124138 (36), prepared in-house). For each RNA construct, a 10 mL reaction mixture containing T7 RNA Polymerase, 0.2-1.0 µM DNA template, transcription buffer (40 mM Tris pH 8.0, 1 mM spermidine, 0.01% Triton-X), 20-50 mM MgCl_2_, 2.5 mM DTT, 10-20% DMSO, and 1 mM each of either unlabeled rATP, rCTP, rUTP, rGTP (MP Biomedicals) or uniformly ^13^C, ^15^N-labeled rATP, rCTP, rUTP, rGTP (Cambridge Isotope Laboratories) was incubated at 37 °C for 6-8 hours. RNA constructs were purified by 10-20% denaturing polyacrylamide gel electrophoresis (PAGE). The RNA band was visualized by UV shadowing with a handheld UV lamp at 254 nm. After band excision, RNA was eluted from the gel using the ‘crush and soak’ method (37) by incubating gel pieces in crush and soak buffer (300 mM sodium acetate pH 5.2, 1 mM EDTA) for 24-48 hours at room temperature with rocking. RNA was further purified to remove acrylamide contaminants by ion-exchange chromatography using a diethylaminoethanol (DEAE) column (GE Healthcare) and elution into buffer (10 mM sodium phosphate pH 7.6, 1 mM EDTA, 1.5 M KCl). RNA was diluted to <100 μM in ultrapure water and annealed by heating to 95 °C for 3 minutes, followed by snap cooling on ice for 1 hour. RNA was then buffer exchanged into the appropriate buffer using a 3-10 kDa Amicon concentrator (Millipore Sigma).

SL3 domain constructs for chemical probing were prepared as previously described (27). Briefly, constructs were designed with flanking 5’ and 3’ overhang sequences for primer annealing during downstream PCR of double stranded DNA (dsDNA) and reverse transcription to complementary DNA (cDNA), respectively. Primers for PCR assembly of the DNA templates were designed using Primerize (38). PCR was performed using Q5 DNA Polymerase (NEB) followed by purification using DNA clean and concentrator spin-column purification (Zymo). RNA was transcribed and purified as described above and stored at −80 °C in ultrapure water until use.

### NMR spectroscopy

Solution NMR spectroscopy experiments were performed at 5 °C, 25 °C, and 40 °C on a Bruker Neo 600 MHz NMR spectrometer equipped with a triple-resonance HCN cryoprobe. NMR samples were prepared in NMR buffer (20 mM sodium phosphate, 50 mM KCl, pH 7.5) with added 5% D_2_O at 0.1-0.8 mM concentrations in 3 mm NMR tubes (Norell). Exchangeable (H1, H3, H41, H42) and nonexchangeable (H2, H5, H6, H8, H1’) proton resonances were assigned using 2D ^1^H-^1^H NOESY spectra of unlabeled RNA samples with mixing times ranging from 100 ms to 250 ms using SD_noe11ezg (39) and noesyesggph pulse sequences. ^1^H-^15^N HSQC and ^1^H-^13^C HSQC spectra were collected using ^13^C/^15^N labeled RNA samples. Data were processed using NMRPipe (40) and analyzed using NMRFAM-Sparky 1.470 powered by Sparky 3.190 (41) in the NMRbox virtual machine (42).

For MgCl_2_ titration experiments, 0.5-0.8 mM RNA samples were titrated with 2, 6 and 10 equivalents of MgCl_2_ and ^1^H-^15^N HSQC and ^1^H-^13^C HSQC spectra were collected at each titration point. Unlabeled RNA samples with 10 equivalents of MgCl_2_ were used to acquire 2D ^1^H–^1^H NOESY spectra at 250 ms for proton resonance assignments and NOE connectivities. Weighted average chemical shift perturbations were calculated using the equation √ΔH^2^ + 0.25Δ*C*^2^ (43).

### DMS-MaPseq

DMS-MaPseq experiments were performed at 25 °C using established protocols (27,44). RNA samples were annealed by heating at 90 °C in water for two minutes then 4 °C for five minutes, followed by addition of DMS-MaPseq folding buffer (100 mM HEPES, pH 8.0, with 60 mM Na^+^ after correcting pH with NaOH) with 0-10 mM MgCl_2_. DMS-MaPseq experiments were performed as described previously (27). Briefly, a 15% DMS (Sigma Aldrich) stock solution in ethanol was used for DMS-MaPseq experiments. 7.5 pmol RNA was incubated with 1.5% DMS solution at 25 °C for six minutes in a 25 μL total reaction volume, then quenched with 25 µL of 100% β-mercaptoethanol (BME). DMS-treated samples were purified using RNA clean and concentrator-5 spin-column kits (Zymo). As a control, reactions were performed using an equivalent volume of ethanol to estimate mutations induced by reverse transcriptase enzyme in the absence of DMS treatment. Each experiment was performed in duplicate.

### Library preparation for sequencing

cDNA was generated using Marathon (Kerafast) reverse transcriptase. RNA was reverse transcribed (RT) using 100 mM DTT, 10 mM dNTPs, RT buffer (250 mM Tris-HCl pH 8.0, 375 mM KCl, 15 mM MgCl_2_), and primers with custom barcodes for downstream demultiplexing. The RT reaction was incubated at 42 °C for three hours. The reaction was quenched by adding 5 µL of 0.4 M NaOH to a 12 µL RT reaction (final concentration of 167 mM NaOH), then heated at 90 °C for 2 minutes and cooled at 4 °C, followed by neutralization with the addition of 2.5 µL quench acid (5 M NaCl, 2 M HCl, 3 M sodium acetate). The cDNA was purified using Zymo oligo clean and concentrator-5 kit (Zymo). To prepare dsDNA for sequencing, purified cDNA was used as a template for PCR using Q5 DNA Polymerase (NEB) and forward and reverse sequencing primers. PCR was performed using 16 cycles with a 15 second extension time and an annealing temperature of 62 °C. PCR products were visualized on a 4% agarose E-gel (Invitrogen) and gel purified using an Invitrogen gel purification kit (Invitrogen) following the manufacturer’s protocol. The dsDNA samples were quantified using a qubit 1x dsDNA high-sensitivity assay kit (Invitrogen) and qubit flex fluorometer (Invitrogen). Individual dsDNA samples of equal fractions were pooled together to generate a 1 nM dsDNA library. 25% PhiX (Illumina) was added to the final library and loaded on the sequencer. RNA constructs were sequenced on the NextSeq 1000 (Illumina) using NextSeq 1000/2000 P1 XLEAP-SBS reagent kit, 2 x 150 read length (Illumina). The total number of reads were 500,000-1,500,000 per sample.

### DMS-MaPseq statistical analyses and visualization

DMS-MaPseq sequencing data was processed using DREEM as previously described (27). To account for the stochastic nature of the DREEM algorithm, three iterations of DREEM clustering were performed for each experimental replicate to calculate the average mutation fraction (MF) values, SL3a and SL3e populations, and standard deviation. Data was visualized using in-house python scripts (https://github.com/eichhorn-lab) that utilize SciPy (45), Pandas (46), Matplotlib (47), NumPy (48), Seaborn (49) and Biopython (50) modules. Final secondary structure models were prepared using VARNA (51). Mutation fraction perturbation (MFP) values were calculated by calculating the difference in the MF value for experiments performed with 0 and 10 mM MgCl_2_. The magnesium midpoint, [Mg^2+^]_1/2_, was calculated using a four-parameter Hill

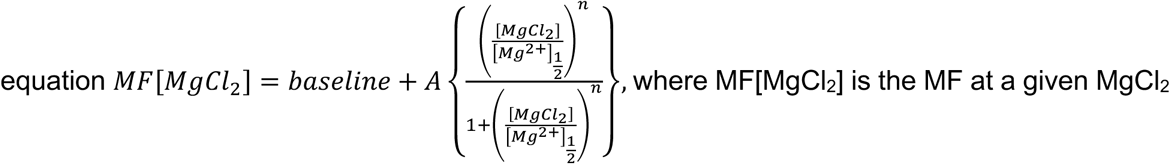

concentration, A is the amplitude, baseline is the MF in the initial state in the absence of MgCl_2_, [Mg^2+^]_1/2_ is MgCl_2_ concentration at the half-maximal change, and n is the Hill coefficient or cooperativity factor. Bootstrap resampling was used to estimate error. If the MFP was less than 0.002 units, the residue was considered non-responsive to MgCl_2_. Experimentally determined populations were used to calculate the free energy difference between SL3a and SL3e states (ΔG_EA_) using the Boltzmann equation ΔG=RTln[K], where K is [SL3e]/[SL3a]. Raw MF values and fitted values are provided in **Supplementary Files 2 and 3**, respectively.

### Isothermal titration calorimetry (ITC)

ITC experiments were performed using a Malvern ITC-200 microcalorimeter (Micro-cal PEAQ-ITC). RNA samples were exchanged in DMS-MaPseq folding buffer. Solid MgCl_2_•6H_2_O (Sigma-Aldrich) was dissolved in DMS-MaPseq folding buffer. For monovalent cation screening, 1 M NaCl was included in the DMS-MaPseq folding buffer. The concentrations of RNA samples in the cell ranged from 10 to 20 µM and MgCl_2_ in the syringe was 500 µM. Titration experiments were performed at 25 °C with 19 injection points, 2 μL for each injection point, with 150 second spacing between injections. Data were fitted to a one-site binding model (11,52) using the Micro-cal PEAQ-ITC analysis software to obtain [Mg^2+^]_1/2_, n, ΔH, and ΔG fitted values. Each experiment was repeated independently in duplicate or triplicate. To determine statistically significant differences in [Mg^2+^]_1/2_ values for RNA constructs, a Welch’s t-test was performed for pairs of RNA constructs (53).

### Linkage equilibrium model

An in-house python script was generated to predict SL3a and SL3e populations from ITC K_D_ values using a generalized linkage equilibrium model (54,55):

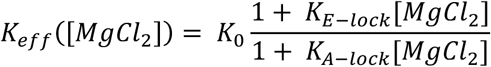

Where 𝐾*_eff_*([𝑀𝑔𝐶𝑙_2_]) is the effective conformational equilibrium constant at a given MgCl_2_ concentration; 𝐾_0_ or ([SL3e]/[SL3a]), is the initial conformational equilibrium constant at low concentrations of MgCl_2_ (0-50 μM), 𝐾*_E-lock_* is the binding constant for E-lock construct, representing the SL3e state; and 𝐾*_A-Lock_* is the binding constant for A-lock construct, representing the SL3a state.

## RESULTS

### Addition of Mg^2+^ shifts SL3 two-state equilibrium to SL3e as the major state due to differential affinities to Mg^2+^

We previously identified that the distal hairpin of human 7SK SL3 exists in a two-state equilibrium, with SL3e and SL3a states having near-equal populations (**Fig. 1A**). To determine the influence of Mg^2+^ on this equilibrium, we performed DMS-MaPseq studies on wild-type (WT) human SL3 with increasing MgCl_2_ concentrations from 0-10 mM at 25 °C. DMS-MaPseq data was clustered using DREEM (27,56) to determine the populations of SL3e and SL3a states at each MgCl_2_ concentration (**Fig. 1B and Supplementary Table S3**). Consistent with our prior studies, we observe a two-state system with SL3a and SL3e conformers at near-equal populations low levels (<100 μM) of MgCl_2_, corresponding to a standard free energy difference between the two states (ΔG°_EA_) of 0.08 ± 0.01 kcal/mol. However, with increasing MgCl_2_ we observed a cooperative transition consistent with redistribution of the two-state equilibrium to SL3e as the major state, corresponding to a ΔG°_EA_ of 0.42 ± 0.006 kcal/mol. These data indicate that addition of Mg^2+^ increases the free energy difference between SL3e and SL3a states by 0.34 ± 0.01 kcal/mol.

We hypothesized that the observed population shift may be due to cation affinity differences for SL3e and SL3a states. To evaluate Mg^2+^ binding to SL3e and SL3a conformers, we used rationally designed RNA constructs that we previously established lock SL3 into either the SL3e state (C224A, E-lock) or SL3a state (ΔC221, A-lock) and performed Mg^2+^ binding measurements using isothermal titration calorimetry (ITC) (**Fig. 1C, Supplementary Fig. 1**). For both constructs, endothermic binding was observed indicating entropic rather than enthalpic contributions to binding (12,57). We used a generalized one-site model to obtain apparent fitting parameters. The apparent Mg^2+^ midpoint concentrations ([Mg^2+^]_1/2_) are in the micromolar range where the [Mg^2+^]_1/2_ of the E-lock construct is approximately 1.7-fold lower than the A-lock construct (**Fig. 1C, Supplementary Table S4**). The corresponding difference in the free energy of binding (ΔΔG_bind_) is 0.33 ± 0.1 kcal/mol which is remarkably consistent with the ΔΔG°_EA_ measured from DMS-MaPseq experiments. To determine whether Mg^2+^ has specific coordination or diffuse electrostatic interactions with SL3 constructs, we performed ITC measurements in the presence of a 1 M NaCl monovalent cation screen (58–63). Under these high monovalent ionic strength conditions, no change in enthalpy was observed for either E-lock or A-lock constructs, indicating that the observed interactions are due to diffuse electrostatic interactions rather than specific Mg^2+^ coordination (**Supplementary Fig. S1**). Together, these data indicate while that Mg^2+^ associates diffusely with both states, a subtle increased affinity of Mg^2+^ to the SL3e state biases the conformational equilibrium.

The excellent agreement in free energy differences for DMS-MaPseq and ITC experiments suggest that preferred association of Mg^2+^ to the SL3e state is minimally sufficient to explain the observed population shift. However, the 1.7-fold difference in [Mg^2+^]_1/2_ between A-lock and E-lock constructs is quite modest. To identify if the affinity difference measured by ITC is sufficient to explain the experimentally determined population shift from DMS-MaPseq experiments, we applied a linkage equilibrium model (54,55) to develop a thermodynamic framework to describe how Mg^2+^ association and SL3 two-state equilibrium are coupled. In this model, we used the [Mg^2+^]_1/2_ measured by ITC of locked constructs to predict the populations of SL3a and SL3e states at low (1 μM) and high (1 mM) MgCl_2_ concentrations. Consistent with experimentally determined populations, the model predicts 54 ± 2% SL3e at 1 μM MgCl_2_ and 66 ± 5% SL3e at 1 mM MgCl_2_ (**Fig. 1D**). From these data, we generated a thermodynamic cycle summarizing these findings (**Fig. 1E**) to illustrate how Mg^2+^ association is thermodynamically linked to the SL3 conformational equilibrium. Together, these data show that subtle differences in affinity can produce quantitative shifts in populations in multi-state RNAs.

### Conformer-locked constructs reveal differential and region-specific Mg^2+^ responses

In our prior study, A-lock and E-lock constructs (**Fig. 2A**) showed excellent agreement to mutation fraction (MF) data of respective WT SL3a and SL3e clusters (27). To further evaluate how well these constructs reflect WT behavior in the presence of divalent cations, we performed DMS-MaPseq experiments with 0 and 10 mM MgCl_2_ and compared MF values between WT SL3a and SL3e clusters and respective locked constructs. We found that the E-lock construct showed excellent agreement to the WT SL3e cluster in both conditions (R^2^ = 0.92 and 0.97, respectively) (**Fig. 2B, Supplementary Fig. S2A-B**). When comparing MF values between 0 and 10 mM MgCl_2_, several residues showed changes in the absolute magnitude, hereafter called the mutation fraction perturbation (MFP). The per-residue MFP comparing the WT SL3e cluster and E-lock construct also showed good agreement (R^2^ = 0.88) (**Fig. 2C**). MFP values for J2/3 residues C229 and A245 are increased in the E-lock construct compared to the WT SL3e cluster, and when excluding these two residues the correlation moderately improves (R^2^ = 0.94). For the A-lock construct, we observe good agreement to the WT SL3a cluster (R^2^ = 0.86), as previously demonstrated (27). Excluding J1/2 loop residues, where the A-lock sequence variation is located (ΔC221), moderately improves the correlation (R^2^ = 0.95). While still strong, the correlation is reduced with 10 mM MgCl_2_ (R^2^ = 0.78), and similarly improves when excluding J1/2 residues (R^2^ = 0.81) (**Fig. 2D, Supplementary Fig. S2C-D**). MFP values also show good agreement between the A-lock construct and SL3a state and improve when excluding J1/2 residues (**Fig. 2E**). Significant MFP values were observed for residues in the upper region for both constructs, particularly the J2/3 loop and apical loop. Consistent with the 2-fold difference in apparent [Mg^2+^]_1/2_ from ITC data, the MFP values showed an approximately 2-fold larger magnitude in the E-lock construct compared to A-lock construct, with 1σ standard deviation values of 0.028 and 0.012, respectively (**Fig. 2F**).

**Figure 2.**
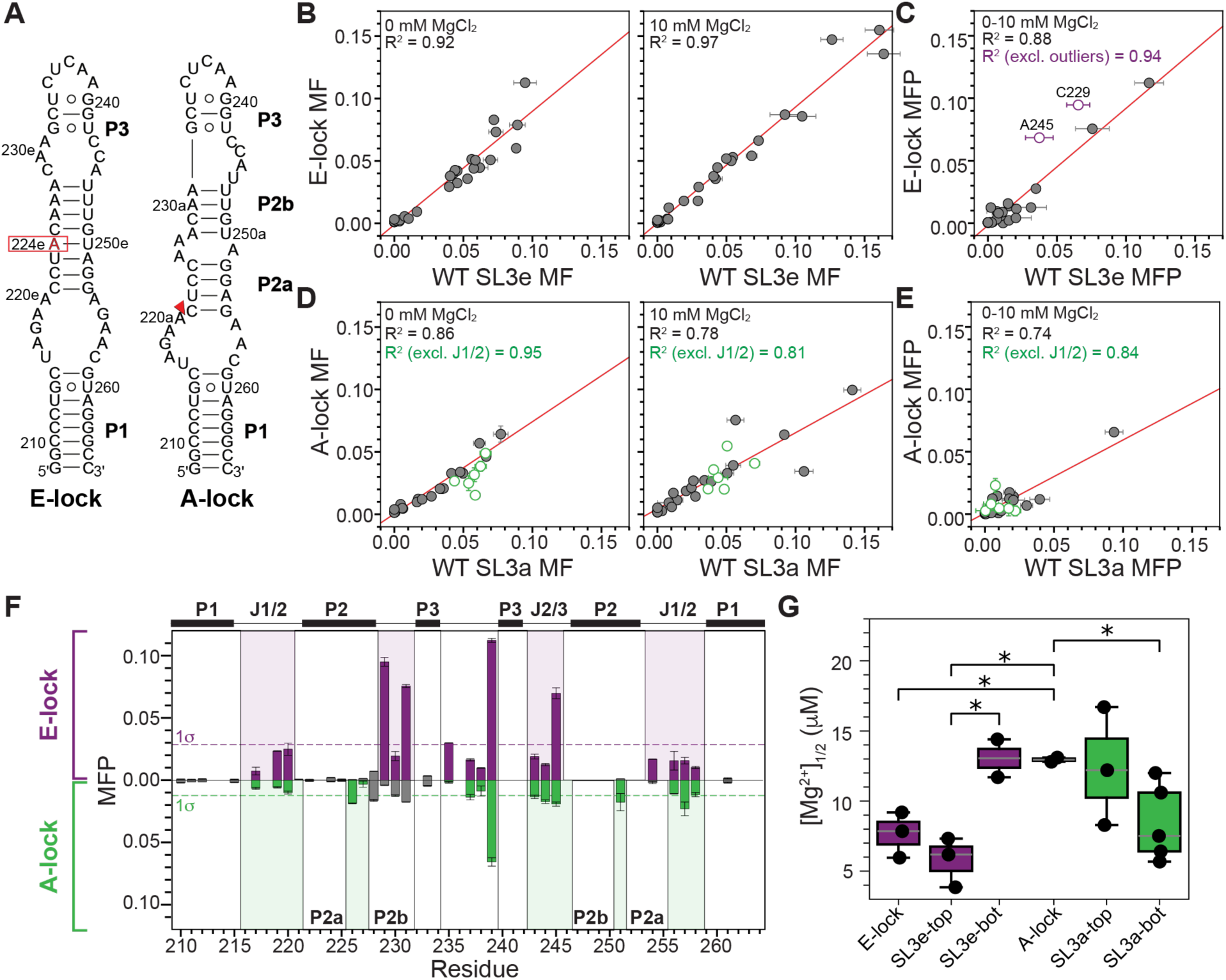
DMS-MaPseq measurements show quantitative agreement for WT clusters and conformer-locked constructs and localize regions of Mg^2+^ association. **(A)** Secondary structures of E-lock and A-lock constructs that stabilize the SL3e and SL3a conformers, respectively. The red box and triangle indicate substitution introduced to lock SL3e state (C224) or SL3a state (ΔC221), respectively. **(B-C)** Correlation between E-lock DMS mutation fraction (MF) and WT SL3e MF at 0 mM MgCl_2_ (**B, left)** and 10 mM MgCl_2_ **(B, right)**, and correlation of mutation fraction perturbations (MFP) between 0 and 10 mM MgCl_2_ data **(C)**. Two outliers (C229 and A245) are labeled. **(D-E)** Analogous correlations for A-lock construct compared to WT SL3a conformer. Green circles indicate J1/2 junction residues that deviate from the correlation. **(F)** SL3 residue-specific MFPs for E-lock (purple) and A-lock (green) constructs. Dashed lines indicate ±1σ thresholds. Structural regions are labeled. **(G)** Magnesium midpoint values ([Mg^2+^]_1/2_) determined from ITC MgCl_2_ titrations for SL3 constructs. Asterisks indicate statistically significant differences (*P* < 0.05).

After establishing that the conformer-locked constructs reasonably reflect SL3a and SL3e conformer behavior in response to Mg^2+^, we next performed MgCl_2_ titration experiments to quantify residue-specific and region-specific differences in Mg^2+^ response between the two states. Both A-lock and E-lock constructs have residues that show cooperative, concentration-dependent changes in MF value in response to MgCl_2_ (**Supplementary Fig. S3 and Table S5**). We were able to fit these data to a modified Hill equation to determine the midpoint Mg^2+^ concentration producing a half-maximal MFP response ([Mg^2+^]_1/2_) (**Supplementary File S3**). We note that DMS-MaPseq experiments report on the ability of N1 or N3 moieties to chemically react with DMS, and the measured [Mg^2+^]_1/2_ values likely do not reflect the true Mg^2+^ binding affinity. Consistent with ITC data, [Mg^2+^]_1/2_ values were generally ∼2-fold lower for E-lock residues compared to A-lock, particularly in the J2/3, P3 stem, and apical loop region (**Supplementary Fig. S3**). While the apparent [Mg^2+^]_1/2_ is higher in qDMS-MaPseq compared to ITC experiments, the ∼2-fold observed difference in the Mg^2+^ midpoint between A-lock and E-lock constructs is consistent between the two methods, supporting the conclusion that SL3e has enhanced responsiveness to Mg^2+^.

We next inspected DMS-MaPseq MgCl_2_ titration data to evaluate whether observed Mg*^2+^* responsiveness is correlated with secondary structural features (**Supplementary Table S5 and Fig. S3**). Here, to reduce noise in the WT SL3 DMS-MaPseq data associated with conformational exchange and clustering, we inspected data from A-lock and E-lock constructs. The structural differences between SL3a and SL3e states result in several SL3 residues that are paired in one conformer and unpaired in the other conformer. This difference in structural environment may impact a given residue’s responsiveness to Mg^2+^. As expected, P1 stem residues are identical in both SL3a and SL3e states and are not responsive to Mg^2+^ for either A- or E-lock constructs (**Supplementary Fig. S3 and Table S5**). Similarly, residues that are in the P2 (C222, C224, C225) and P3 (C233) stems in both conformers either show no response or slight decrease in the MF value. While the apical loop secondary structure is the same in both states, minor differences are observed in MF values, particularly for C235 and A239 adjacent to the P3 stem. These differences may be due to subtle differences in thermodynamic stability of the P3 stem between SL3a and SL3e states, which can propagate to the apical loop. Generally, residues located in the P2 stem in one state but a loop in the other state, for example C229, is not responsive to Mg^2+^ in the A-lock construct but shows a significant decrease in the MF value in E-lock construct. These data indicate that Mg*^2+^*addition does not significantly impact the reactivity of structured residues with DMS. In contrast, residues located in J1/2 or J2/3 loops in both states showed varying responsiveness to Mg^2+^, likely due to differences in loop structure between the two states and response to Mg^2+^ addition. Taken together, qDMS-MaPseq results and comparative analysis of A-lock and E-lock constructs are consistent with ITC data showing that Mg^2+^ has stronger association with the SL3e state compared to the SL3a state.

From qDMS-MaPseq data, we hypothesized that that upper region of the SL3e state is the primary site for divalent cation association. To test this hypothesis, we used constructs corresponding to top and bottom fragments of SL3a and SL3e states to perform Mg^2+^ binding measurements using ITC (**Fig. 2G, Supplementary Fig. S4**). The SL3e-top fragment and the E-lock construct have comparable [Mg^2+^]_1/2_ values and increased affinity compared to SL3e-bottom, A-lock, and SL3a-state fragment constructs. To evaluate whether measured [Mg^2+^]_1/2_ differences are statistically significant, we performed a Welch’s t-test between constructs and found that the [Mg^2+^]_1/2_ value for the SL3e-top construct is significantly increased compared to either SL3e-bottom or A-lock constructs (p-values 0.0451 and 0.0180, respectively). Both the SL3a-top and - bottom constructs had reduced [Mg^2+^]_1/2_ values compared to SL3e-top. The slightly increased Mg^2+^ affinity of SL3a-bottom compared to A-lock may be due to the C221 deletion in the A-lock construct, which is present in the WT sequence. Consistent with this observation, qDMS-MaPseq MgCl_2_ titration experiments show that C221 has reduced MF values with added MgCl_2_ (**Supplementary Fig. S2**), suggesting Mg^2+^ addition protects C221 from DMS reactivity. The ITC results confirm that Mg^2+^ primarily interacts with the upper part of SL3e, consistent with DMS-MaPseq experiments.

### Identification of an A•C mismatch signature from qDMS-MaPseq data

While inspecting qDMS mutation fraction data from the MgCl_2_ titration experiments, we observed residues in base-paired regions in P2 and P3 stems adjacent to the J2/3 loop including A228 (P2) and C233 (P3) show reduced MF values (**Fig. 3A**), indicating increased solvent protection consistent with increased ordering. When examining adenines opposite cytosines in the J2/3 and apical loops, we observed a recurring pattern in which the cytosines showed a gradual reduced value, indicating protection from DMS methylation, and the adenine showed increased values, indicating greater DMS methylation. These patterns were observed for J2/3 residues C229-A245 and A231-C243, as well as apical loop residues C235-A239 (**Fig. 3A and Supplementary Fig. S3**). Notably, J2/3 residues A230-C244 do not show this pattern. Rather, both A230 and C244 show modest increases in MFP, indicating both residues have increased solvent accessibility. This pattern suggests a signature of potential A•C wobble base pairs that is consistent with a hydrogen bonding scheme where the C N3 is participating in a hydrogen bond but the A N1 is solvent exposed (**Fig. 3B-C**).

**Figure 3.**
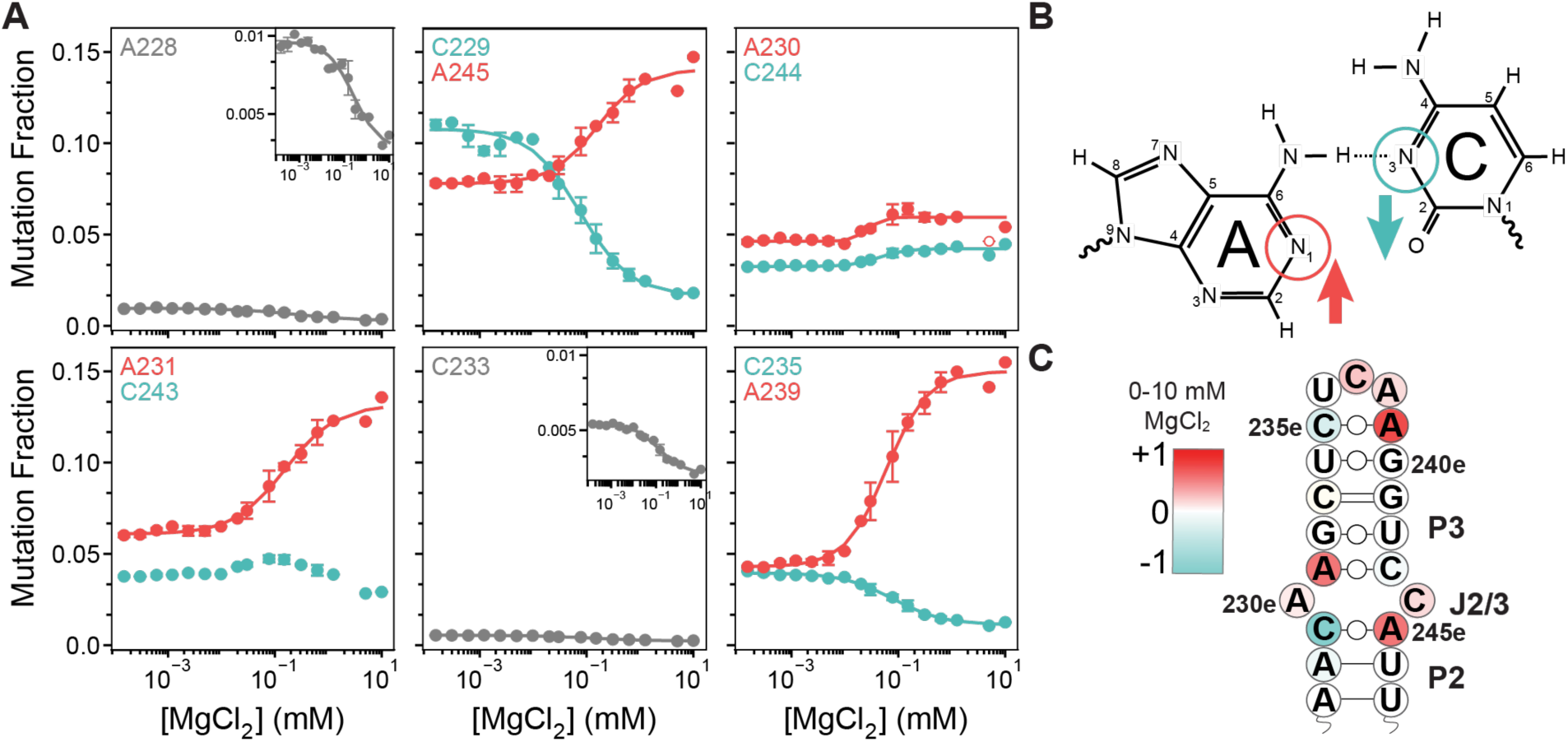
DMS reactivity patterns reveal A•C wobble base pair signatures from MgCl_2_ titration experiments. **(A)** Mutation fraction values of DMS-MaPseq MgCl_2_ titration experiments for nucleotides in the J2/3 loop and P3 stem. Residues are color-coded by structural context (red: adenines in J2/3 loop; cyan: cytosines in J2/3 loop; gray: A228 at the terminal P2 base pair adjacent to J2/3 loop, C233 in the P3 stem). **(B)** Chemical structure of proposed A•C wobble base pair with Watson-Crick face DMS modification sites circled. Arrows indicate direction of MFP values with increasing MgCl_2_. **(C)** Secondary structure of the SL3e upper region color-coded by DMS MFP values, where red indicates increased MF values and cyan indicates decreased MF values.

### Addition of Mg^2+^ stabilizes G•U-rich P3 stem

Both DMS-MaPseq and ITC experiments indicate Mg^2+^ association to the SL3e upper region. To obtain atomic-level insights into cation-mediated impacts to this region, we performed MgCl_2_ titration experiments using solution NMR spectroscopy for the SL3e-top construct at pH 7.5 to maintain similar conditions with DMS-MaPseq experiments (**Fig. 4A)**. ^1^H-^15^N HSQC experiments were performed to monitor exchangeable imino N1H1 and N3H3 resonances. Imino protons for P3 stem residues (U232, U234, G240, G241, U242) are not observable at pH 7.5, even at 5 °C, likely due to solvent exchange (**Fig. 4B).** However, upon addition of MgCl_2_ all P3 imino resonances appear and increase in intensity with increasing MgCl_2_ (**Fig. 4B-C**). Minimal weighted-average chemical shift perturbations (CSPs) are observed, indicating that Mg^2+^ association results in reduced solvent exchange of imino protons rather than changes to the chemical environment.

**Figure 4.**
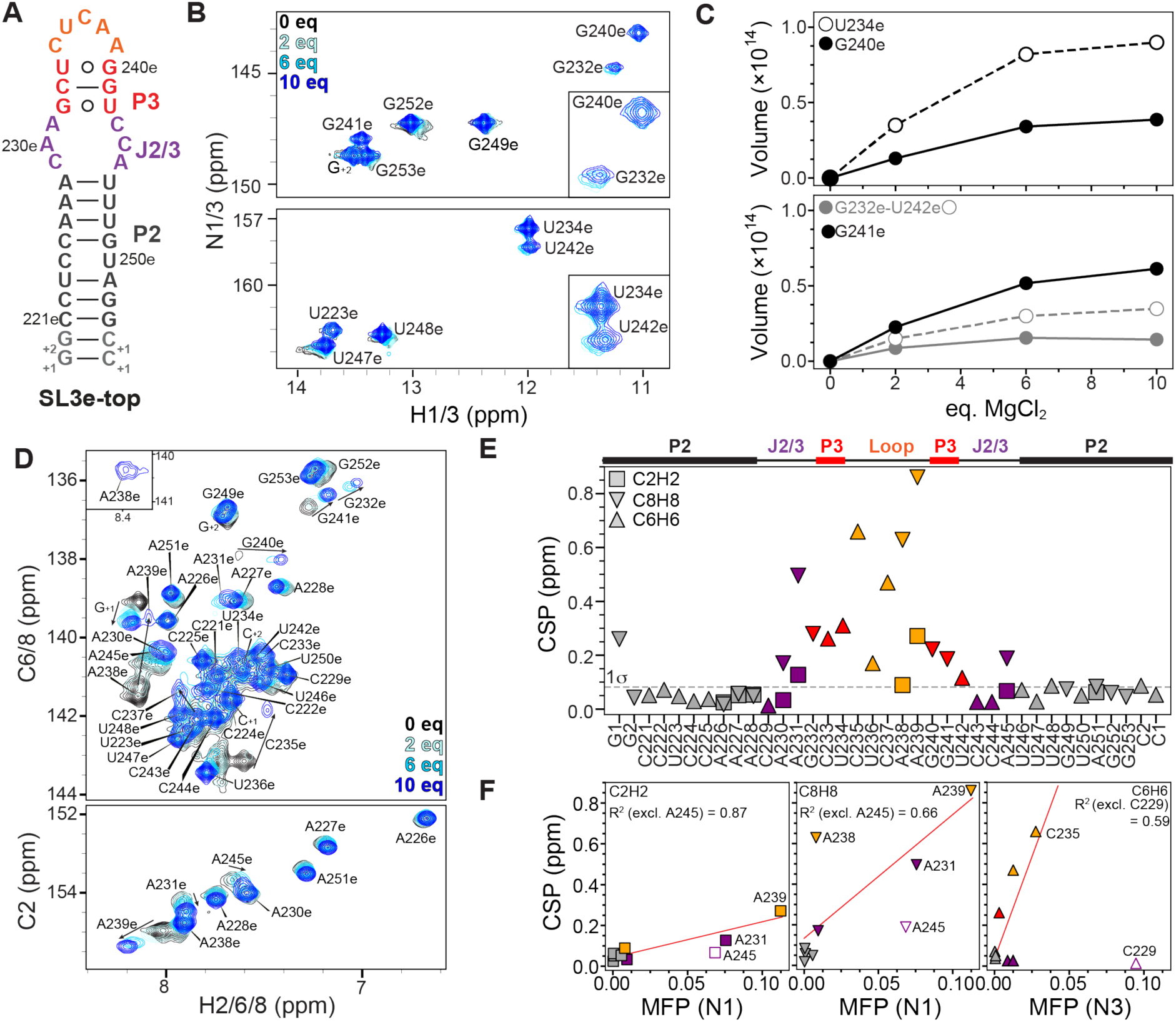
NMR spectroscopy reveals Mg^2+^ stabilizes the SL3e-state, and Mg^2+^-dependent chemical shift perturbations are correlated with DMS reactivity. **(A)** Secondary structure of the SL3e-top construct used for NMR experiments colored by structural region: gray, P2 stem; purple, J2/3 loop; red, P3 stem; orange, apical loop. **(B)** ^1^H-^15^N HSQC spectrum of imino resonances show minimal chemical shift perturbations (CSPs) at 0 eq (black), 2 eq (ice blue), 6 eq (cyan), and 10 eq (blue) MgCl_2_. Inset panels highlight G232, G240, U234, and U242 resonances in G•U wobble pairs. The ‘e’ subscript denotes SL3e conformer assignments **(C)** MgCl_2_ titration data showing peak volume changes for P3 stem imino resonances. **(D)** ^1^H-^13^C HSQC spectrum of aromatic resonances, colored by MgCl_2_ eq as indicated in panel **B**. **(E)** CSPs for 0 and 10 eq MgCl_2_ for C2H2 (squares), C8H8 (triangles), and C6H6 (inverted triangles) resonances. **(F)** Correlation plots between NMR CSPs and DMS MFPs for C2H2, C8H8, and C6H6 to N1, N1, and N3 respectively. R^2^ values are indicated. Outliers (A245 and C229) are labeled.

We performed ^1^H-^13^C HSQC experiments to monitor nucleobase nonexchangeable C2H2, C6H6, C8H8 resonances (**Fig. 4D and Supplementary Fig. S5**). We observed substantial CSPs in the J2/3 region, P3 stem, and apical loop residues (**Fig. 4E and Supplementary Fig. S6**). The P2 stem showed minimal CSPs, with the exception of the terminal G_+1_ residue. Together, the ITC, DMS-MaPseq, and NMR data support a model where cations diffusely interact with the P3 stem and apical loop to increase ordering of the P3 stem. While Mg^2+^-induced DMS MFPs are observed in the J2/3 loop (**Fig. 3A**), NMR MgCl_2_ titrations show significant CSPs only for P3 and apical loop residues, suggesting that Mg^2+^ stabilization of the P3 stem indirectly results in ordering of the J2/3 loop through enhanced coaxial stacking interactions.

We next examined if NMR CSPs were correlated to DMS-MaPseq MFPs (**Fig. 4F**). NMR chemical shift reports on local electronic environment, while DMS mutation fraction reports on ability to chemically react with DMS. The adenine C2H2, located on the Watson-Crick face adjacent to the N1 position, has excellent agreement for all residues (R^2^ = 0.72) with improved agreement when excluding A245 (R^2^ = 0.87), an above-identified outlier. Adenine C8H8 and cytosine C6H6 show reasonable agreement (R^2^ = 0.66 and 0.59, respectively) after excluding A245 and C229, which were previously identified as outliers. These residues show larger MFP values relative to CSP. In contrast, A238 C8H8 has a larger CSP than MFP, suggesting addition of Mg^2+^ changes the electronic environment of this moiety to a greater extent than changes to solvent accessibility. The correlation in CSP and MFP indicates that both are reporting on the same effect of Mg^2+^-induced structural stabilization of the SL3e upper stem.

### Mg^2+^ promotes an A-form like geometry in the SL3e upper stem

After establishing that Mg^2+^ primarily associates with the upper stem region of SL3e, we next examined the molecular details of Mg^2+^-induced structural changes in the SL3e-top fragment construct using 2D ^1^H-^1^H NOESY (nuclear Overhauser effect spectroscopy). We performed complete sequential assignment through NOESY walks of H1’, H2, H5, H6, and H8 proton cross-strand connectivities at both 25°C and 40 °C in the absence and presence of 10 mM MgCl_2_ (**Supplementary Fig. S7-S9**). At 25 °C, significant resonance overlap was observed for protons including U250 H6, U246 H6, C229 H6, A228 H8, C221 H6, and G_+2_ H8 **(Supplementary Fig. S7)**. Increasing the temperature to 40 °C improved spectral resolution **(Supplementary Fig. S9 A-B)**. As expected, P2 stem residues showed NOE connectivities that were consistent with an A-form helical geometry in both the presence and absence of Mg^2+^ **(Supplementary Fig. S7-S9).** Addition of Mg^2+^ resulted in the appearance of numerous new and well-resolved NOEs, particularly in the P3 stem region (**Fig. 5A-C, Supplementary Fig. S10**). Addition of Mg^2+^ enabled a continuous NOE walk from the P2 stem, J2/3 loop, and P3 stem regions, consistent with an A-form-like helical structure in the J2/3 loop and P3 stem (**Fig. 5D, Supplementary Figs. S8-S9**). In the absence of Mg^2+^, the J2/3 loop exhibited limited NOE connectivities, with only sparse H1’–H8 and H2–H1’ NOEs among adenine and cytosine residues, suggesting partial base stacking (**Supplementary Fig. S7**). Upon addition of MgCl_2_, the number and intensity of NOE cross-peaks increased substantially, including both sugar-to-base and base-to-base interactions throughout the J2/3 loop (**Supplementary Figs. S8–S9**). In the absence of Mg^2+^, base-to-base NOEs were observed only from A239 H8 through to C243 H6 (**Supplementary Fig. S7)**. In contrast in the presence of 10 equivalents of Mg^2+^, base-to-base NOE connectivities extended from C229 H6 in the J2/3 loop through the P3 stem to U236 H6, with missing NOEs in the apical loop between U236-A238, then from A238 H8 to U247 H6 **(Supplementary Fig. S8)**. This pattern indicates Mg^2+^-stabilized base stacking and adoption of an ordered, A-form-like helical geometry in the symmetric J2/3 loop. Together, NMR and DMS-MaPseq data support a model where the symmetric J2/3 loop has an A-form like geometry in the presence of MgCl_2_, with at least two A•C wobble base pairs.

**Figure 5.**
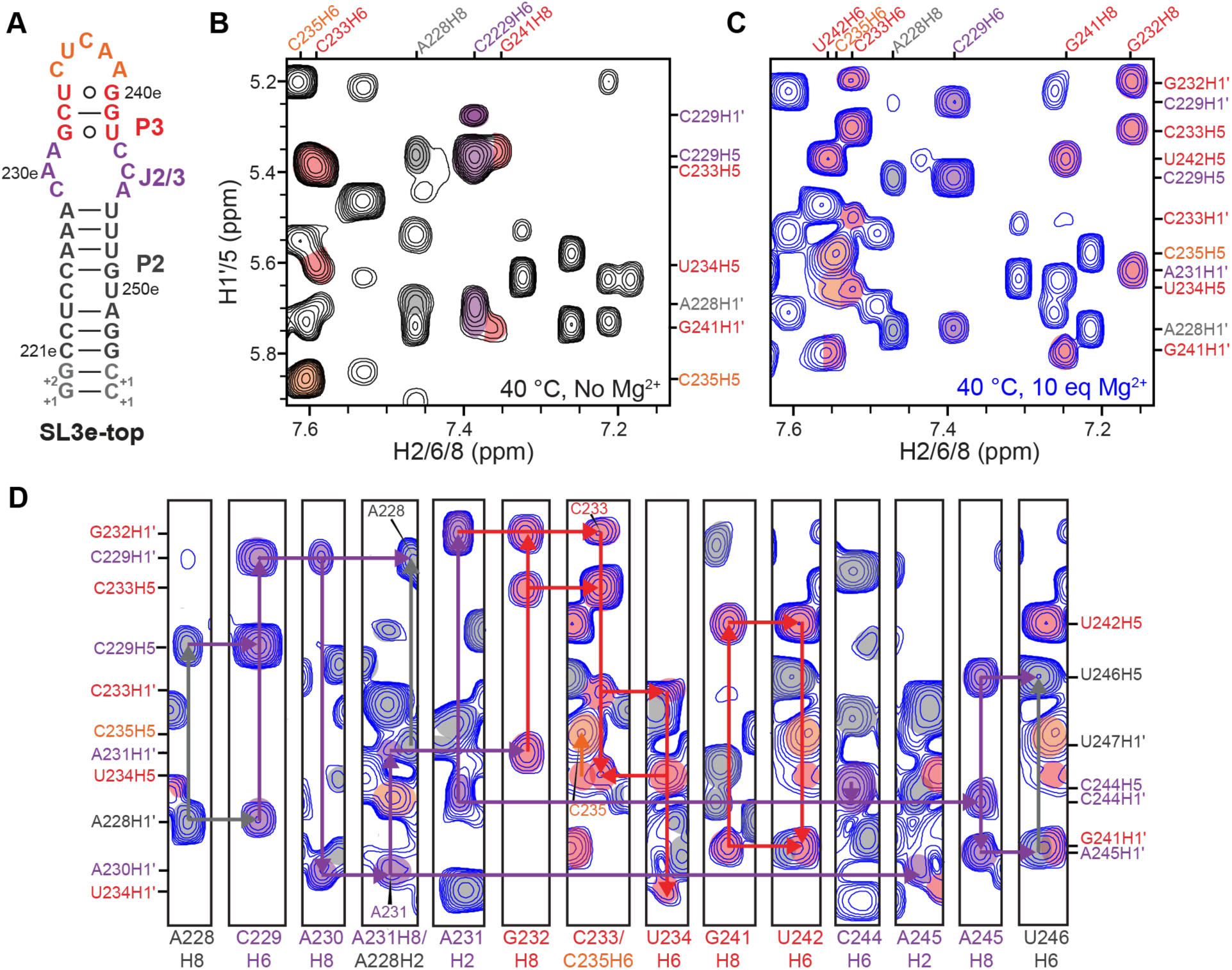
Addition of Mg^2+^ reveals new and well resolved NOEs and an A-form-like geometry in the SL3e upper stem. **(A)** Secondary structure of the SL3e-top construct used for NMR experiments. **(B-C)** ^1^H-^1^H NOESY spectral region showing the different NOEs for the P3 stem **(B)** in the absence of Mg^2+^ and **(C)** in the presence of Mg^2+^. **(D)** Sequential walk of ^1^H-^1^H NOESY spectrum of SL3e-top construct with 10 eq MgCl_2_ showing NOE connectivity patterns in the J2/3 region and P3 stem. Gray lines indicate NOE connectivities for P2 stem, purple lines indicate NOE connectivities for J2/3 loop, red lines indicate NOE connectivities for P3 stem, and orange line indicate NOE connectivities for the apical loop.

## DISCUSSION

It is increasingly recognized that RNA exists as an ensemble of conformational states rather than a single, static structure. However, the fundamental thermodynamic principles that define population distributions within RNA ensembles, and how changing environmental conditions modulate this equilibrium, are not fully characterized. Here, we integrate qDMS-MaPseq, NMR spectroscopy, and ITC experiments to dissect the Mg^2+^-dependent equilibrium of the 7SK SL3 domain. A central finding of this study is that a relatively small free energy difference in Mg^2+^ association between SL3 states (ΔΔG ∼0.3 kcal/mol) produces a measurable shift in population (**Fig. 6A**). While such energetic differences are often considered minimal in the context of RNA folding, our results demonstrate that they are sufficient to redistribute populations within a multi-state ensemble. The strong agreement between ΔΔG values derived from DMS-MaPseq population analysis and ITC binding measurements provides support for a linkage equilibrium model in which Mg^2+^ association is thermodynamically coupled to population. Importantly, this agreement shows that preferential Mg^2+^ association to the SL3e state is minimally sufficient to explain the observed population shift. While the data are well described by a two-state model in both the absence and presence of Mg^2+^, we cannot exclude the potential that lowly populated states may be present that are not resolved by the methods used in this study. Nevertheless, the consistency across independent approaches indicates that the two-state model captures the dominant features of this system. The differential Mg^2+^ affinity between SL3e and SL3a states can be explained by distinct structural differences, particularly in loop motifs. In SL3e, the J2/3 loop adopts a more continuously stacked geometry that pre-organizes the RNA for P3 stem formation, reducing the entropic cost of folding. In contrast, the J2/3 loop in the SL3a state contains an asymmetric bulge that disrupts P2-P3 stem stacking and introduces conformational flexibility, increasing the entropic cost of folding. Mg^2+^ preferentially associates with the more pre-organized SL3e state, shifting the SL3 two-state equilibrium through a conformational selection-type mechanism.

**Figure 6.**
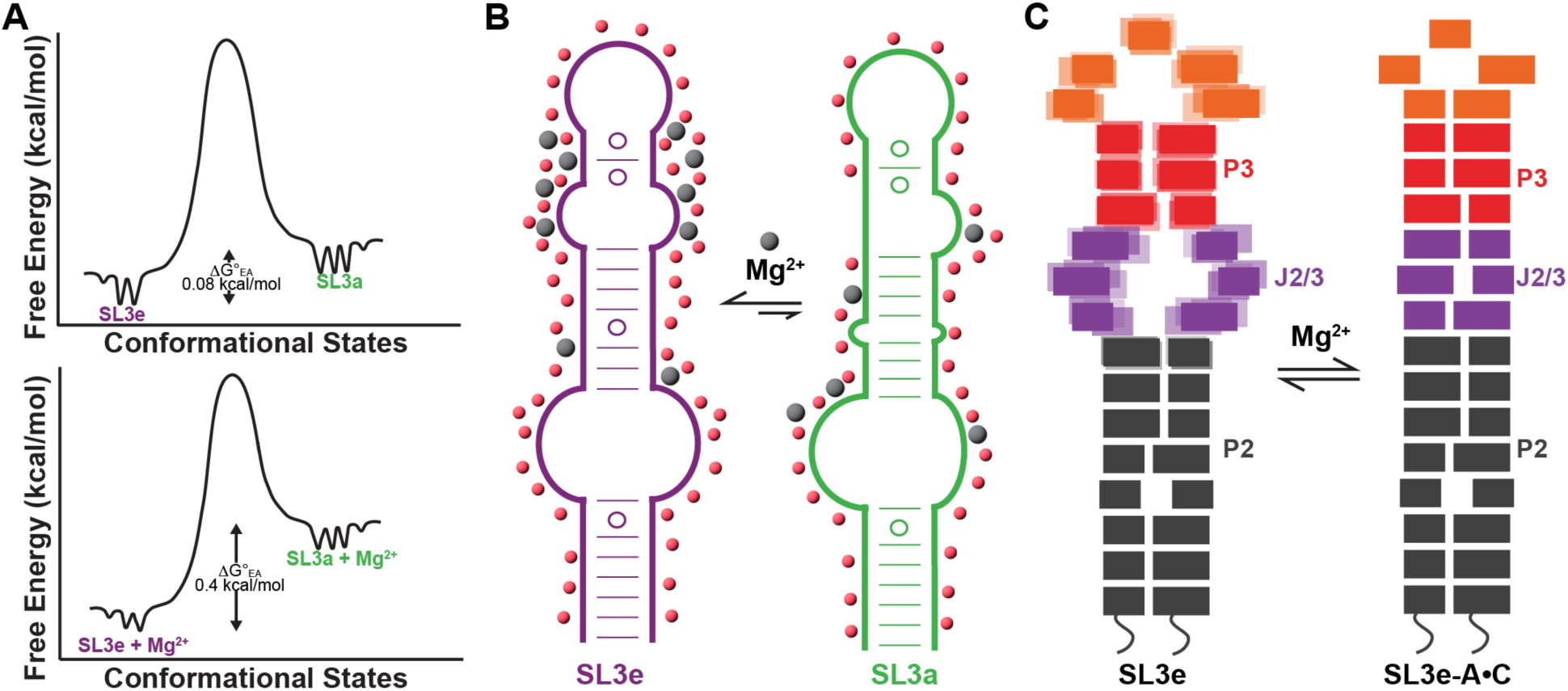
Model of Mg^2+^-driven redistribution of 7SK SL3 conformational ensemble. **(A)** Free energy conformational landscape to visualize SL3e and SL3a equilibrium in the (*top*) absence and (*bottom*) presence of Mg^2+^. **(B**) Cartoon models representing the diffuse association and distribution of Mg^2+^ that shifts SL3 two-state equilibrium toward the SL3e state. Red spheres indicate monovalent cations and gray spheres indicate divalent Mg^2+^ cations. **(C)** Cartoon models representing Mg^2+^ induced structural changes and stabilization in the upper region of the SL3e state.

DMS-MaPseq and ITC experiments of conformer-locked SL3 constructs localize Mg^2+^ association to the upper stem region of the SL3e state, particularly the P3 stem and J2/3 loop (**Fig. 6B**). While sequence substitutions used to lock individual conformers may alter local RNA structural dynamics or RNA-ion interactions, the strong agreement between A-lock and E-lock constructs to their respective WT states indicates that these effects are minimal in this study, though they remain an important consideration when applying this approach to other RNAs. NMR experiments show Mg^2+^-dependent stabilization of the P3 stem, which further reinforces J2/3 loop ordering and promotes coaxial stacking between P2 and P3 stems (**Fig. 6C**). Notably, minimal CSPs of the imino resonances and increased NOEs in the J2/3 loop and P3 stem region suggest that Mg^2+^ acts primarily by stabilizing pre-existing structural features rather than inducing a new structural state. These observations are consistent with nearest-neighbor effects where local structural features influence the stability of adjacent elements (64), extended here to an RNA undergoing conformational exchange. Together, these data support a model in which Mg^2+^ promotes a more ordered, A-form-like geometry in the upper stem of the SL3e state (**Fig. 6C**).

While divalent cations such as Mg^2+^ are well-established to stabilize individual RNA structures and promote tertiary folding (10,12,13), less is known about how cations redistribute populations among RNA with alternative folded states. In this study, Mg^2+^ interactions with SL3 are predominantly diffuse rather than site-specific; however, these interactions are not uniformly distributed across the RNA but are localized to the upper region of the SL3e state. In addition to a pre-organized SL3e J2/3 loop and P3 stem, the P3 stem contains two G•U wobble base pairs, which are known to associate with Mg^2+^ (65). However, the endothermic enthalpy change observed in ITC MgCl_2_ titration experiments indicates that Mg^2+^ association is entropically rather than enthalpically driven, consistent with diffuse electrostatic interactions involving charge screening and release of water molecules (12,60). Consistent with this interpretation ITC experiments performed under high monovalent salt conditions, which compete with diffuse Mg^2+^ association, show no detectable change in enthalpy, further supporting a nonspecific association model.

DMS-MaPseq has been used quantitatively to measure [Mg^2+^]_1/2_ values for RNA tertiary contacts and structural motifs (21,22). However, qDMS-MaPseq has not been demonstrated for RNAs that adopt multiple conformations. A notable outcome of this work is the strong quantitative agreement between Mg^2+^-induced perturbations measured by DMS-MaPseq and NMR spectroscopy. The strong correlation between DMS MFPs and NMR CSPs further validates that DMS reactivity can serve as a sensitive reporter of local structural changes. The strongest correlation was observed between DMS MFP and NMR CSP for respective adenine N1 and C2H2 moieties, as both the DMS-reactive N1 position and the C2H2 moiety are located on the Watson-Crick face. Although less strong, moderate correlations were also observed for adenine N1 and C8H8 and cytosine N3 and C6H6 moieties. Future studies employing chemical probing techniques that directly probe the Hoogsteen edge N7 position (64,65) may further improve the correlation with C8H8 CSPs. While chemical probing approaches are often interpreted qualitatively, our results demonstrate that DMS-MaPseq can provide quantitative, residue-level insight into conformational equilibria when combined with appropriate analysis frameworks.

In addition to providing insight into SL3 conformational equilibria, our data reveal a characteristic DMS reactivity signature for A•C wobble base pairs. Identifying and characterizing noncanonical base pairs remains challenging without solving a high-resolution structure. To study these noncanonical base pairs, approaches such as ion- and pH-dependent NMR studies, thermodynamic analysis, and chemical probing are increasingly utilized (66–69). Here, we observe a reproducible pattern in DMS-MaPseq MgCl_2_ titration experiments in which cytosine residues exhibit Mg^2+^ dose-dependent decreased MF values while the cross-strand adenines show increased MF values. This behavior is consistent with a wobble base pairing geometry in which the cytosine N3 participates in hydrogen bonding while the adenine N1 remains solvent exposed (**Fig. 3B**). Given the challenges associated with identifying noncanonical base pairs, particularly A•C wobbles, this signature may provide a useful indicator for detecting such interactions in other RNAs. While further validation is required, these findings expand the utility of chemical probing for identifying noncanonical structural features.

Although this study is primarily focused on thermodynamic and structural characterization, our findings have potential implications for the biological function of 7SK RNA. The SL3 domain serves as a critical hub for protein recruitment within the 7SK ribonucleoprotein complex (34,35), and population shifts arising from changes in cellular ionic strength could influence the accessibility of protein-binding surfaces. In this context, ion-dependent modulation of RNA ensembles may represent a mechanism for regulating RNP assembly, analogous to assembly of the ribosome (70,71) and riboswitch aptamer-ligand complexes (72–75). Future studies incorporating protein binding and additional cellular conditions will be necessary to directly test these hypotheses. In summary, this work demonstrates that modest differences in Mg^2+^ association can quantitatively reshape RNA conformational ensembles and establishes an integrated experimental and thermodynamic framework for analyzing these effects. By combining DMS-MaPseq, NMR spectroscopy, and calorimetry, we provide an approach for linking ion binding to structural and energetic features of RNA ensembles. While additional studies will be required to determine how general this behavior is across other RNAs and cellular conditions, these results provide a framework for investigating how ionic environments modulate RNA structure and dynamics.

## DATA AVAILABILITY

Python scripts are deposited in GitHub (https://github.com/eichhorn-lab). Raw demultiplexed chemical probing data have been deposited in the NCBI BioProject (XXXX). Processed chemical probing data is deposited in GitHub (https://github.com/eichhorn-lab). NMR resonance assignments for SL3e-top in the absence and presence of MgCl_2_ have been deposited to the BMRB (ID XXXX).

## SUPPLEMENTARY DATA

The following supplementary data are available: **Supplementary File 1**, which contains supplemental tables and figures; **Supplementary File 2**, which contains raw DMS-MaPseq mutation fraction values; and **Supplementary File 3**, which contains qDMS-MaPseq fitted values from the MgCl_2_ titration data.

## AUTHOR CONTRIBUTIONS

CDE conceived and oversaw all aspects of the project. MBC, SOA, and CDE designed experiments and performed data analysis. MBC and SOA prepared samples and performed experiments. MBC, SOA, and CDE wrote the paper with input from all authors.

## ACKNOWLEDGEMENTS

We acknowledge Dr. Joseph D. Yesselman, Sakshi Jain, and Gavin Anderson for assistance with sequencing. We thank Dr. Martha Morton for NMR assistance. We acknowledge Addgene for the pQE30-His-T7RNAP plasmid used to prepare T7 RNA Polymerase enzyme for *in vitro* transcribed RNA samples, which was a gift from Sebastian Maerkl & Takuya Ueda (Addgene plasmid #124138).

## REFERENCES

1. Cech, T.R. and Steitz, J.A. (2014) The noncoding RNA revolution-trashing old rules to forge new ones. Cell, 157, 77–94.

2. Al-Hashimi, H.M. and Walter, N.G. (2008) RNA dynamics: it is about time. Curr Opin Struct Biol, 18, 321–329.

3. Ganser, L.R., Kelly, M.L., Herschlag, D. and Al-Hashimi, H.M. (2019) The roles of structural dynamics in the cellular functions of RNAs. Nat Rev Mol Cell Biol, 20, 474–489.

4. Bonilla, S.L., Jones, A.N. and Incarnato, D. (2024) Structural and biophysical dissection of RNA conformational ensembles. Curr Opin Struct Biol, 88, 102908.

5. Dethoff, E.A., Petzold, K., Chugh, J., Casiano-Negroni, A. and Al-Hashimi, H.M. (2012) Visualizing transient low-populated structures of RNA. Nature, 491, 724–728.

6. Mustoe, A.M., Brooks, C.L. and Al-Hashimi, H.M. (2014) Hierarchy of RNA functional dynamics. Annu Rev Biochem, 83, 441–466.

7. Baisden, J.T., Boyer, J.A., Zhao, B., Hammond, S.M. and Zhang, Q. (2021) Visualizing a protonated RNA state that modulates microRNA-21 maturation. Nat Chem Biol, 17, 80–88.

8. Leontis, N.B., Lescoute, A. and Westhof, E. (2006) The building blocks and motifs of RNA architecture. Curr Opin Struct Biol, 16, 279–287.

9. Guth-Metzler, R., Mohamed, A.M., Cowan, E.T., Henning, A., Ito, C., Frenkel-Pinter, M., Wartell, R.M., Glass, J.B. and Williams, L.D. (2023) Goldilocks and RNA: where Mg2+ concentration is just right. Nucleic Acids Res, 51, 3529–3539.

10. Romani, A.M. (2011) Cellular magnesium homeostasis. Arch Biochem Biophys, 512, 1–23.

11. Laing, L.G., Gluick, T.C. and Draper, D.E. (1994) Stabilization of RNA structure by Mg ions. Specific and non-specific effects. J Mol Biol, 237, 577–587.

12. Draper, D.E. (2004) A guide to ions and RNA structure. RNA, 10, 335–343.

13. Draper, D.E. (2008) RNA folding: thermodynamic and molecular descriptions of the roles of ions. Biophys J, 95, 5489–5495.

14. Xu, D., Landon, T., Greenbaum, N.L. and Fenley, M.O. (2007) The electrostatic characteristics of G.U wobble base pairs. Nucleic Acids Res, 35, 3836–3847.

15. Stefan, L.R., Zhang, R., Levitan, A.G., Hendrix, D.K., Brenner, S.E. and Holbrook, S.R. (2006) MeRNA: a database of metal ion binding sites in RNA structures. Nucleic Acids Res, 34, D131–134.

16. Zheng, H., Shabalin, I.G., Handing, K.B., Bujnicki, J.M. and Minor, W. (2015) Magnesium-binding architectures in RNA crystal structures: validation, binding preferences, classification and motif detection. Nucleic Acids Res, 43, 3789–3801.

17. Kim, H.D., Nienhaus, G.U., Ha, T., Orr, J.W., Williamson, J.R. and Chu, S. (2002) Mg2+-dependent conformational change of RNA studied by fluorescence correlation and FRET on immobilized single molecules. Proc Natl Acad Sci U S A, 99, 4284–4289.

18. Wu, M. and Tinoco, I., Jr. (1998) RNA folding causes secondary structure rearrangement. Proc Natl Acad Sci U S A, 95, 11555–11560.

19. Xue, Y., Gracia, B., Herschlag, D., Russell, R. and Al-Hashimi, H.M. (2016) Visualizing the formation of an RNA folding intermediate through a fast highly modular secondary structure switch. Nat Commun, 7, ncomms11768.

20. Weidmann, C.A., Mustoe, A.M., Jariwala, P.B., Calabrese, J.M. and Weeks, K.M. (2021) Analysis of RNA-protein networks with RNP-MaP defines functional hubs on RNA. Nat Biotechnol, 39, 347–356.

21. Lange, B., Gil, R.G., Anderson, G.S. and Yesselman, J.D. (2024) High-throughput determination of RNA tertiary contact thermodynamics by quantitative DMS chemical mapping. Nucleic Acids Res, 52, 9953–9965.

22. Deenalattha, D.H.S., Jurich, C.P., Lange, B., Armstrong, D., Nein, K. and Yesselman, J.D. (2025) Characterizing 3D RNA structural features from DMS reactivity. bioRxiv.

23. Olson, S.W., Turner, A.W., Arney, J.W., Saleem, I., Weidmann, C.A., Margolis, D.M., Weeks, K.M. and Mustoe, A.M. (2022) Discovery of a large-scale, cell-state-responsive allosteric switch in the 7SK RNA using DANCE-MaP. Mol Cell, 82, 1708–1723 e1710.

24. Aviran, S. and Incarnato, D. (2022) Computational Approaches for RNA Structure Ensemble Deconvolution from Structure Probing Data. J Mol Biol, 434, 167635.

25. Tomezsko, P.J., Corbin, V.D.A., Gupta, P., Swaminathan, H., Glasgow, M., Persad, S., Edwards, M.D., McIntosh, L., Papenfuss, A.T., Emery, A. et al. (2020) Determination of RNA structural diversity and its role in HIV-1 RNA splicing. Nature, 582, 438–442.

26. Gherghe, C.M., Shajani, Z., Wilkinson, K.A., Varani, G. and Weeks, K.M. (2008) Strong correlation between SHAPE chemistry and the generalized NMR order parameter (S2) in RNA. J Am Chem Soc, 130, 12244–12245.

27. Camara, M.B., Lange, B., Yesselman, J.D. and Eichhorn, C.D. (2024) Visualizing a two-state conformational ensemble in stem-loop 3 of the transcriptional regulator 7SK RNA. Nucleic Acids Res, 52, 940–952.

28. Barboric, M., Lenasi, T., Chen, H., Johansen, E.B., Guo, S. and Peterlin, B.M. (2009) 7SK snRNP/P-TEFb couples transcription elongation with alternative splicing and is essential for vertebrate development. Proc Natl Acad Sci U S A, 106, 7798–7803.

29. Brogie, J.E. and Price, D.H. (2017) Reconstitution of a functional 7SK snRNP. Nucleic Acids Res, 45, 6864–6880.

30. Nguyen, V.T., Kiss, T., Michels, A.A. and Bensaude, O. (2001) 7SK small nuclear RNA binds to and inhibits the activity of CDK9/cyclin T complexes. Nature, 414, 322–325.

31. Yang, Z., Zhu, Q., Luo, K. and Zhou, Q. (2001) The 7SK small nuclear RNA inhibits the CDK9/cyclin T1 kinase to control transcription. Nature, 414, 317–322.

32. Marz, M., Donath, A., Verstraete, N., Nguyen, V.T., Stadler, P.F. and Bensaude, O. (2009) Evolution of 7SK RNA and its protein partners in metazoa. Mol Biol Evol, 26, 2821–2830.

33. Camara, M.B., Sobeh, A.M. and Eichhorn, C.D. (2023) Progress in 7SK ribonucleoprotein structural biology. Front Mol Biosci, 10, 1154622.

34. Van Herreweghe, E., Egloff, S., Goiffon, I., Jady, B.E., Froment, C., Monsarrat, B. and Kiss, T. (2007) Dynamic remodelling of human 7SK snRNP controls the nuclear level of active P-TEFb. EMBO J, 26, 3570–3580.

35. Briese, M., Saal-Bauernschubert, L., Ji, C., Moradi, M., Ghanawi, H., Uhl, M., Appenzeller, S., Backofen, R. and Sendtner, M. (2018) hnRNP R and its main interactor, the noncoding RNA 7SK, coregulate the axonal transcriptome of motoneurons. Proc Natl Acad Sci U S A, 115, E2859–E2868.

36. Shimizu, Y., Inoue, A., Tomari, Y., Suzuki, T., Yokogawa, T., Nishikawa, K. and Ueda, T. (2001) Cell-free translation reconstituted with purified components. Nat Biotechnol, 19, 751–755.

37. Green, M.R. and Sambrook, J. (2019) Isolation of DNA Fragments from Polyacrylamide Gels by the Crush and Soak Method. Cold Spring Harbor Protocols, 2019, pdb.prot100479.

38. Tian, S., Yesselman, J.D., Cordero, P. and Das, R. (2015) Primerize: automated primer assembly for transcribing non-coding RNA domains. Nucleic Acids Res, 43, W522–526.

39. Macaya, R.F., Schultze, P., Smith, F.W., Roe, J.A. and Feigon, J. (1993) Thrombin-binding DNA aptamer forms a unimolecular quadruplex structure in solution. Proc Natl Acad Sci U S A, 90, 3745–3749.

40. Delaglio, F., Grzesiek, S., Vuister, G.W., Zhu, G., Pfeifer, J. and Bax, A. (1995) NMRPipe: a multidimensional spectral processing system based on UNIX pipes. J Biomol NMR, 6, 277–293.

41. Lee, W., Tonelli, M. and Markley, J.L. (2015) NMRFAM-SPARKY: enhanced software for biomolecular NMR spectroscopy. Bioinformatics, 31, 1325–1327.

42. Maciejewski, M.W., Schuyler, A.D., Gryk, M.R., Moraru, II, Romero, P.R., Ulrich, E.L., Eghbalnia, H.R., Livny, M., Delaglio, F. and Hoch, J.C. (2017) NMRbox: A Resource for Biomolecular NMR Computation. Biophys J, 112, 1529–1534.

43. Cavanagh, J. (2007) Protein NMR spectroscopy : principles and practice. 2nd ed. Academic Press, Amsterdam; Boston.

44. Cheng, C.Y., Kladwang, W., Yesselman, J.D. and Das, R. (2017) RNA structure inference through chemical mapping after accidental or intentional mutations. Proc Natl Acad Sci U S A, 114, 9876–9881.

45. Virtanen, P., Gommers, R., Oliphant, T.E., Haberland, M., Reddy, T., Cournapeau, D., Burovski, E., Peterson, P., Weckesser, W., Bright, J. et al. (2020) SciPy 1.0: fundamental algorithms for scientific computing in Python. Nat Methods, 17, 261–272.

46. McKinney, W. (2010) Data structures for statistical computing in python. Proceedings of the 9th Python in Science Conference, 445.

47. Hunter, J.D. (2007) Matplotlib: A 2D Graphics Environment. Computing in Science & Engineering, 9, 90–95.

48. Harris, C.R., Millman, K.J., van der Walt, S.J., Gommers, R., Virtanen, P., Cournapeau, D., Wieser, E., Taylor, J., Berg, S., Smith, N.J., et al. (2020) Array programming with NumPy. Nature, 585, 357–362.

49. Waskom, M., Botvinnik, O., O’Kane, D., Hobson, P., Lukauskas, S., Gemperline, D., Augspurger, T., Halchenko, Y., Cole, J., Warmenhoven, J., et al. (2017) mwaskom/seaborn: v0.8.1 (September 2017). Zenodo.

50. Cock, P.J., Antao, T., Chang, J.T., Chapman, B.A., Cox, C.J., Dalke, A., Friedberg, I., Hamelryck, T., Kauff, F., Wilczynski, B. et al. (2009) Biopython: freely available Python tools for computational molecular biology and bioinformatics. Bioinformatics, 25, 1422–1423.

51. Darty, K., Denise, A. and Ponty, Y. (2009) VARNA: Interactive drawing and editing of the RNA secondary structure. Bioinformatics, 25, 1974–1975.

52. Misra, V.K. and Draper, D.E. (2001) A thermodynamic framework for Mg2+ binding to RNA. Proc Natl Acad Sci U S A, 98, 12456–12461.

53. West, R.M. (2021) Best practice in statistics: Use the Welch t-test when testing the difference between two groups. Ann Clin Biochem, 58, 267–269.

54. Wyman, J. and Gill, S.J. (1990) Binding and linkage : functional chemistry of biological macromolecules. University Science Books, Mill Valley, Ca.

55. Wyman, J. (1967) Allosteric Linkage. Journal of the American Chemical Society, 89, 2202-&.

56. Tomezsko, P., Swaminathan, H. and Rouskin, S. (2021) DMS-MaPseq for Genome-Wide or Targeted RNA Structure Probing In Vitro and In Vivo. Methods Mol Biol, 2254, 219–238.

57. Johnson, C.M. (2021) Isothermal Titration Calorimetry. Methods Mol Biol, 2263, 135–159.

58. Takamoto, K., Das, R., He, Q., Doniach, S., Brenowitz, M., Herschlag, D. and Chance, M.R. (2004) Principles of RNA compaction: insights from the equilibrium folding pathway of the P4-P6 RNA domain in monovalent cations. J Mol Biol, 343, 1195–1206.

59. Lipfert, J., Doniach, S., Das, R. and Herschlag, D. (2014) Understanding nucleic acid-ion interactions. Annu Rev Biochem, 83, 813–841.

60. Das, R., Travers, K.J., Bai, Y. and Herschlag, D. (2005) Determining the Mg2+ stoichiometry for folding an RNA metal ion core. J Am Chem Soc, 127, 8272–8273.

61. Bukhman, Y.V. and Draper, D.E. (1997) Affinities and selectivities of divalent cation binding sites within an RNA tertiary structure. J Mol Biol, 273, 1020–1031.

62. Wilson, T.J. and Lilley, D.M. (2002) Metal ion binding and the folding of the hairpin ribozyme. RNA, 8, 587–600.

63. Record, M.T., Jr., Lohman, M.L. and De Haseth, P. (1976) Ion effects on ligand-nucleic acid interactions. J Mol Biol, 107, 145–158.

64. Xia, T., SantaLucia, J., Jr., Burkard, M.E., Kierzek, R., Schroeder, S.J., Jiao, X., Cox, C. and Turner, D.H. (1998) Thermodynamic parameters for an expanded nearest-neighbor model for formation of RNA duplexes with Watson-Crick base pairs. Biochemistry, 37, 14719–14735.

65. Auffinger, P., Ennifar, E. and D’Ascenzo, L. (2021) Deflating the RNA Mg(2+) bubble: stereochemistry to the rescue! RNA, 27, 243–252.

66. Kotar, A., Ma, S. and Keane, S.C. (2022) pH dependence of C*A, G*A and A*A mismatches in the stem of precursor microRNA-31. Biophys Chem, 283, 106763.

67. Wilcox, J.L. and Bevilacqua, P.C. (2013) pKa shifting in double-stranded RNA is highly dependent upon nearest neighbors and bulge positioning. Biochemistry, 52, 7470–7476.

68. Allawi, H.T. and SantaLucia, J., Jr. (1998) Nearest-neighbor thermodynamics of internal A.C mismatches in DNA: sequence dependence and pH effects. Biochemistry, 37, 9435–9444.

69. Faison, E.M., Nallathambi, A. and Zhang, Q. (2023) Characterizing Protonation-Coupled Conformational Ensembles in RNA via pH-Differential Mutational Profiling with DMS Probing. J Am Chem Soc, 145, 18773–18777.

70. Abeysirigunawardena, S.C. and Woodson, S.A. (2015) Differential effects of ribosomal proteins and Mg2+ ions on a conformational switch during 30S ribosome 5’-domain assembly. RNA, 21, 1859–1865.

71. Wang, A., Levi, M., Mohanty, U. and Whitford, P.C. (2022) Diffuse Ions Coordinate Dynamics in a Ribonucleoprotein Assembly. J Am Chem Soc, 144, 9510–9522.

72. Noeske, J., Schwalbe, H. and Wohnert, J. (2007) Metal-ion binding and metal-ion induced folding of the adenine-sensing riboswitch aptamer domain. Nucleic Acids Res, 35, 5262–5273.

73. Reining, A., Nozinovic, S., Schlepckow, K., Buhr, F., Furtig, B. and Schwalbe, H. (2013) Three-state mechanism couples ligand and temperature sensing in riboswitches. Nature, 499, 355–359.

74. Choudhary, P.K. and Sigel, R.K. (2014) Mg(2+)-induced conformational changes in the btuB riboswitch from E. coli. RNA, 20, 36–45.

75. Latham, M.P., Zimmermann, G.R. and Pardi, A. (2009) NMR chemical exchange as a probe for ligand-binding kinetics in a theophylline-binding RNA aptamer. J Am Chem Soc, 131, 5052–5053.

