## Supplementary File 1 for "Quantifying Mg^2+^ dependence on conformational equilibrium in the two-state 7SK RNA stem-loop 3"

### List of supplementary tables and figures:

**Table S1:** DNA primer and template sequences used in DMS-MaPseq studies
**Table S2:** DNA template sequences used in NMR and ITC studies
**Table S3:** Summary of determined population for SL3 WT $MgCl_2$ titration from DMS-MaPseq clustering analysis
**Table S4:** ITC thermodynamic parameters from $MgCl_2$ titration experiments
**Table S5:** Summary of observed changes in DMS-MaPseq $MgCl_2$ titration experiments
**Figure S1:** Representative $MgCl_2$ titration ITC thermograms of SL3 E-lock and SL3 A-lock constructs
**Figure S2:** VARNA plots of mutation fraction perturbation (MFP) values at 0-10 mM $MgCl_2$ for SL3 WT clusters and conformer-locked constructs
**Figure S3:** Comparison of $MgCl_2$-dependent changes in DMS-MaPseq MF values for A-lock and E-lock constructs
**Figure S4**: Representative $MgCl_2$ titration ITC thermograms of SL3 fragment constructs
**Figure S5:** $^1H$-$^{15}N$ and $^1H$-$^{13}C$ HSQC spectra of SL3e-top $MgCl_2$ titration
**Figure S6:** $^1H$-$^{13}C$ HSQC spectra of SL3e-top at 25 °C
**Figure S7**: $^1H$-$^1H$ NOESY spectra of SL3e-top in the absence of $MgCl_2$ at 25 ºC
**Figure S8**: $^1H$-$^1H$ NOESY spectra of SL3e-top with 10 equivalents of $MgCl_2$ at 25 ºC
**Figure S9**: $^1H$-$^1H$ NOESY spectra of SL3e-top in the absence and presence of $MgCl_2$ at 40 ºC.
**Figure S10**: $^1H$-$^1H$ NOESY spectral regions of SL3e-top supporting stabilized P3 stem with $Mg^{2+}$

**Table S1**. DNA primer and template sequences used in DMS-MaPseq studies. Underlined residues correspond to the bacteriophage T7 RNAP promoter sequence for *in vitro* transcription. Highlighted residues are additional 5' and 3' sequences for barcode and adaptor addition.

| DNA template and primer sequences for DMS-MaPseq | |
|---|---|
| SL3 WT DNA | CTAATACGACTCACTATAGGAAGATCGAGTAGATCAAAGGCCCTGCTAGAACCTCCAAACAAGCTCTCAAGGTCCATTTGTAGGAGAACGTAGGGCCAAAGAAACAACAACAACAAC |
| SL3 WT 1F primer | CTAATACGACTCACTATAGGAAGATCGAGTAGATCAAAGGCCCTGC |
| SL3 WT 2R primer | TGAGAGCTTGTTTGGAGGTTCTAGCAGGGCCTTTGATCTACT |
| SL3 WT 3F primer | ACCTCCAAACAAGCTCTCAAGGTCCATTTGTAGGAGAACGTAGGGCCAAAGAAA |
| SL3 WT 4R primer | GTTGTTGTTGTTGTTTCTTTGGCCCTACGTTCTCCTACAAATG |
| SL3 E-lock DNA | CTAATACGACTCACTATAGGAAGATCGAGTAGATCAAAGGCCCTGCTAGAACCTACAAACAAGCTCTCAAGGTCCATTTGTAGGAGAACGTAGGGCCAAAGAAACAACAACAACAAC |
| SL3 E-lock 1F primer | CTAATACGACTCACTATAGGAAGATCGAGTAGATCAAAGGCCCTGC |
| SL3 E-lock 2R primer | GACCTTGAGAGCTTGTTTGTAGGTTCTAGCAGGGCCTTTGATCTACT |
| SL3 E-lock 3F primer | ACAAACAAGCTCTCAAGGTCCATTTGTAGGAGAACGTAGGGCCAA |
| SL3 E-lock 4R primer | GTTGTTGTTGTTGTTTCTTTGGCCCTACGTTCTCCTACAAATG |
| SL3 A-lock DNA | CTAATACGACTCACTATAGGAAGATCGAGTAGATCAAAGGCCCTGCTAGAACTCCAAACAAGCTCTCAAGGTCCATTTGTAGGAGAACGTAGGGCCAAAGAAACAACAACAACAAC |
| SL3 A-lock 1F primer | CTAATACGACTCACTATAGGAAGATCGAGTAGATCAAAGGCCCTGC |
| SL3 A-lock 2R primer | CCTTGAGAGCTTGTTTGGAGTTCTAGCAGGGCCTTTGATCTACT |
| SL3 A-lock 3F primer | CTCCAAACAAGCTCTCAAGGTCCATTTGTAGGAGAACGTAGGGCCAAAGAAA |
| SL3 A-lock 4R primer | GTTGTTGTTGTTGTTTCTTTGGCCCTACGTTCTCCTACAAATG |

**Table S2**. DNA template sequences used in NMR and ITC studies. Underlined residues correspond to the bacteriophage T7 RNAP promoter sequence for *in vitro* transcription.

| **DNA template sequences for NMR and ITC studies** | |
|---|---|
| **SL3 E-state** | |
| SL3 E-lock | CTAATACGACTCACTATAGGCCCTGCTAGAACCTACAAACAAGCTCTCAAGGTCCATTTGTAGGAGAACGTAGGGCC |
| SL3e-top | CTAATACGACTCACTATAGGCCTCCAAACAAGCTCTCAAGGTCCATTTGTAGGCC |
| SL3e-bottom with cUUCGg tetraloop | CTAATACGACTCACTATAGGCCCTGCTACAACCTCTTCGGAGGACAACGTAGGGCC |
| **SL3 A-state** | |
| SL3 A-lock | CTAATACGACTCACTATAGGCCCTGCTAGAACTCCAAACAAGCTCTCAAGGTCCATTTGTAGGAGAACGTAGGGCC |
| SL3a-top | CTAATACGACTCACTATAGGCTCCAAACAAGCTCTCAAGGTCCATTTGTAGGAGCC |
| SL3a-bottom with cUUCGg tetraloop | CTAATACGACTCACTATAGGCCCTGCTAGAACCTCCTTCGGGAGAACGTAGGGCC |

**Table S3**. Summary of determined population for SL3 WT $MgCl_2$ titration from DMS-MaPseq clustering analysis.

| $MgCl_2$ (mM) | SL3e | SL3a | error |
|---|---|---|---|
| 0 | 56 | 44 | 0.50 |
| 0.00045 | 53 | 47 | 1.2 |
| 0.00075 | 51 | 49 | 0.20 |
| 0.0010 | 53 | 47 | 2.0 |
| 0.0030 | 53 | 47 | 1.2 |
| 0.0090 | 54 | 46 | 1.5 |
| 0.015 | 51 | 49 | 0.20 |
| 0.050 | 55 | 45 | 4.0 |
| 0.10 | 62 | 38 | 0.50 |
| 0.2 | 65 | 35 | 0.2 |
| 0.25 | 66 | 34 | 1.6 |
| 0.5 | 67 | 33 | 1.1 |
| 0.75 | 64 | 36 | 0.10 |
| 1.0 | 67 | 33 | 1.6 |
| 1.5 | 67 | 33 | 0.0 |
| 2.5 | 66 | 34 | 2.9 |
| 5.0 | 68 | 32 | 3.7 |
| 10 | 67 | 33 | 0.20 |

**Table S4.** ITC thermodynamic parameters from $MgCl_2$ titration experiments.

| RNA constructs | N (sites) | $K_D$ (mM) | $\Delta H$ (kcal/mol) | $\Delta G_{bind}$ (kcal/mol) | $\Delta S$, at 25 ºC (cal/mol) | Number of replicates |
|---|---|---|---|---|---|---|
| **SL3 E-state** | | | | | | |
| **SL3 E-lock** | 1.13 ± 0.13 | 7.67 ± 1.62 | 1.95 ± 0.10 | -6.99 ± 0.13 | 29.97 ± 0.75 | 3 |
| **SL3e-top** | 0.96 ± 0.22 | 5.78 ± 2.45 | 2.37 ± 0.68 | -7.17 ± 0.27 | 32.01 ± 1.38 | 3 |
| **SL3e-bottom** | 1.14 ± 0.01 | 13.05 ± 1.91 | 2.68 ± 0.12 | -6.67 ± 0.08 | 31.34 ± 0.12 | 2 |
| **SL3 A-state** | | | | | | |
| **SL3 A-lock** | 1.41 ± 0.14 | 12.95 ± 0.21 | 2.43 ± 0.10 | -6.67 ± 0.01 | 30.50 ± 0.31 | 2 |
| **SL3a-top** | 1.04 ± 0.22 | 12.40 ± 4.21 | 2.84 ± 0.60 | -6.72 ± 0.21 | 32.08 ± 1.41 | 3 |
| **SL3a-bottom** | 0.87 ± 0.08 | 8.44 ± 2.74 | 2.53 ± 0.46 | -6.95 ± 0.19 | 31.80 ± 0.94 | 5 |

**Table S5.** Summary of observed changes in DMS-MaPseq $MgCl_2$ titration experiments

| Region | Residues | Response to $Mg^{2+}$ | Correlation |
|---|---|---|---|
| P1 stem | C10-C215, A261 | No response | n/a |
| J1/2 loop in both A and E state | A217, A219, A220 | No response | n/a |
| | A257 | Both increase | Correlated |
| | C258 | E state no response<br>A state decreases | Uncorrelated |
| J1/2 in E only<br>P2a stem in A state | A254 | E state decreases<br>A state no response | Uncorrelated |
| J1/2 in A only<br>P2 stem in E state | C221 | E state no response<br>WT SL3a state decreases | Uncorrelated |
| P2 stem in both | C222, C224/A224, C225 | No response | n/a |
| | A228 | E state decreases<br>A state increases | Anti-correlated |
| A-loop<br>P2 stem in E state | A226 | E state no response<br>A state decreases | Uncorrelated |
| | A227, A251 | No response | n/a |
| J2/3 in E state<br>P2b stem in A state | C229 | E state decreases<br>A state no response | Uncorrelated |
| | A230 | Both increase | Correlated |
| | A231 | Both increase | Correlated |
| P3 stem | C233 | Both decrease | Correlated |
| Apical loop | C235 | E state decreases<br>A state no response | Uncorrelated |
| | C237 | Both increase | Correlated |
| | A238 | E state increases<br>A state no response | Uncorrelated |
| | A239 | Both increase | Correlated |
| J2/3 in both | C243 | E state cannot be fit<br>A state increases | Uncorrelated |
| | C244 | Both increase | Correlated |
| | A245 | E state increases<br>A state decreases | Uncorrelated |

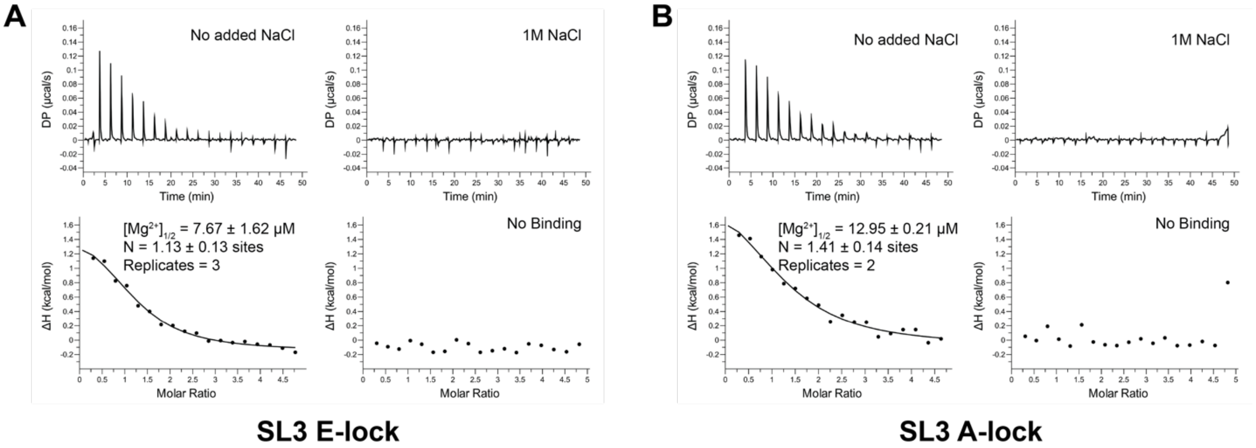


**Figure S1:** Representative $MgCl_2$ titration ITC thermograms of SL3 E-lock and SL3 A-lock constructs. The ITC experiments were performed at 25 °C in a buffer of 100mM HEPES, pH 8.0, and 60 mM NaOH for pH adjustment.

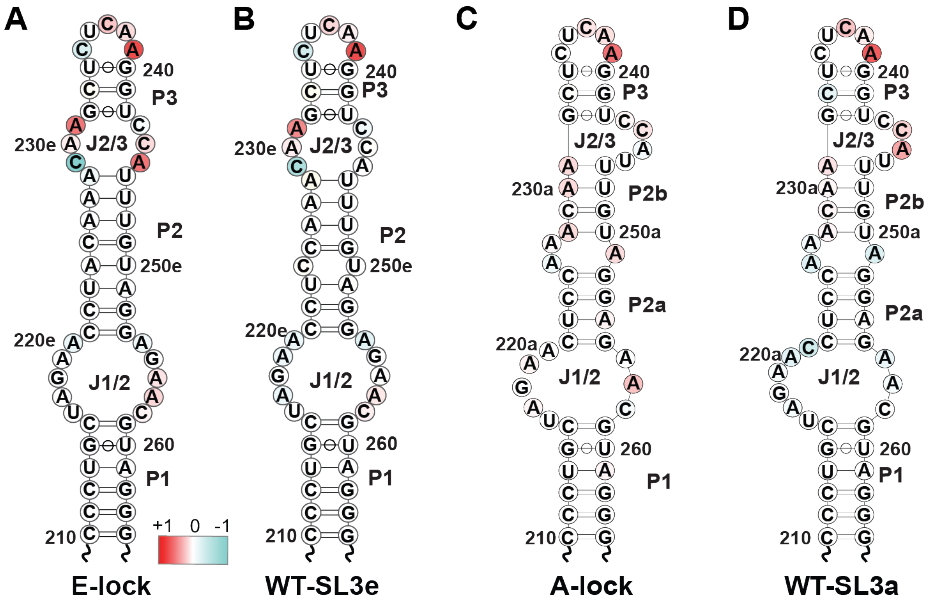


**Figure S2:** VARNA plots of mutation fraction perturbation (MFP) values at 0-10 mM $MgCl_2$ for SL3 WT clusters and conformer-locked constructs. A) E-lock construct, B) WT SL3e cluster, C) A-lock construct, and D) WT SL3a cluster. Residues that have reduced MF values in the presence of $Mg^{2+}$ will have a negative value and are colored blue. Residues that have increased MF values in the presence of $Mg^{2+}$ will have a positive value and are colored red. DMS-MaPseq MF values show the largest changes in the SL3 upper stem.

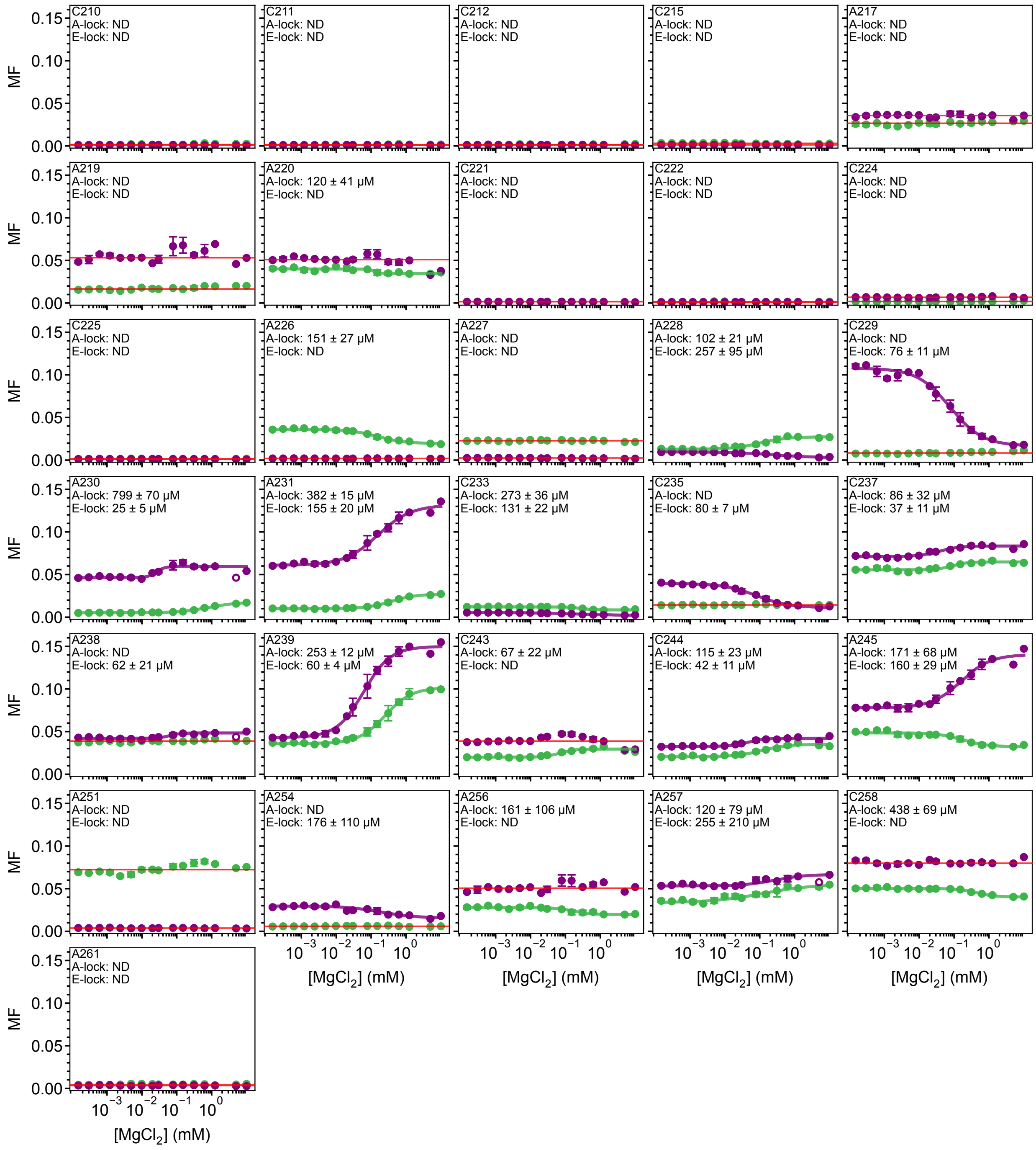


**Figure S3**: Comparison of $MgCl_2$-dependent changes in DMS-MaPseq MF values for A-lock and E-lock constructs. A-lock data is shown in green, and E-lock data is shown in purple. Residues that did not show $MgCl_2$-dependent changes in MF values are depicted with a red line. ND indicates 'not determined'. See **Table 5** for summary of observed changes in the plots.

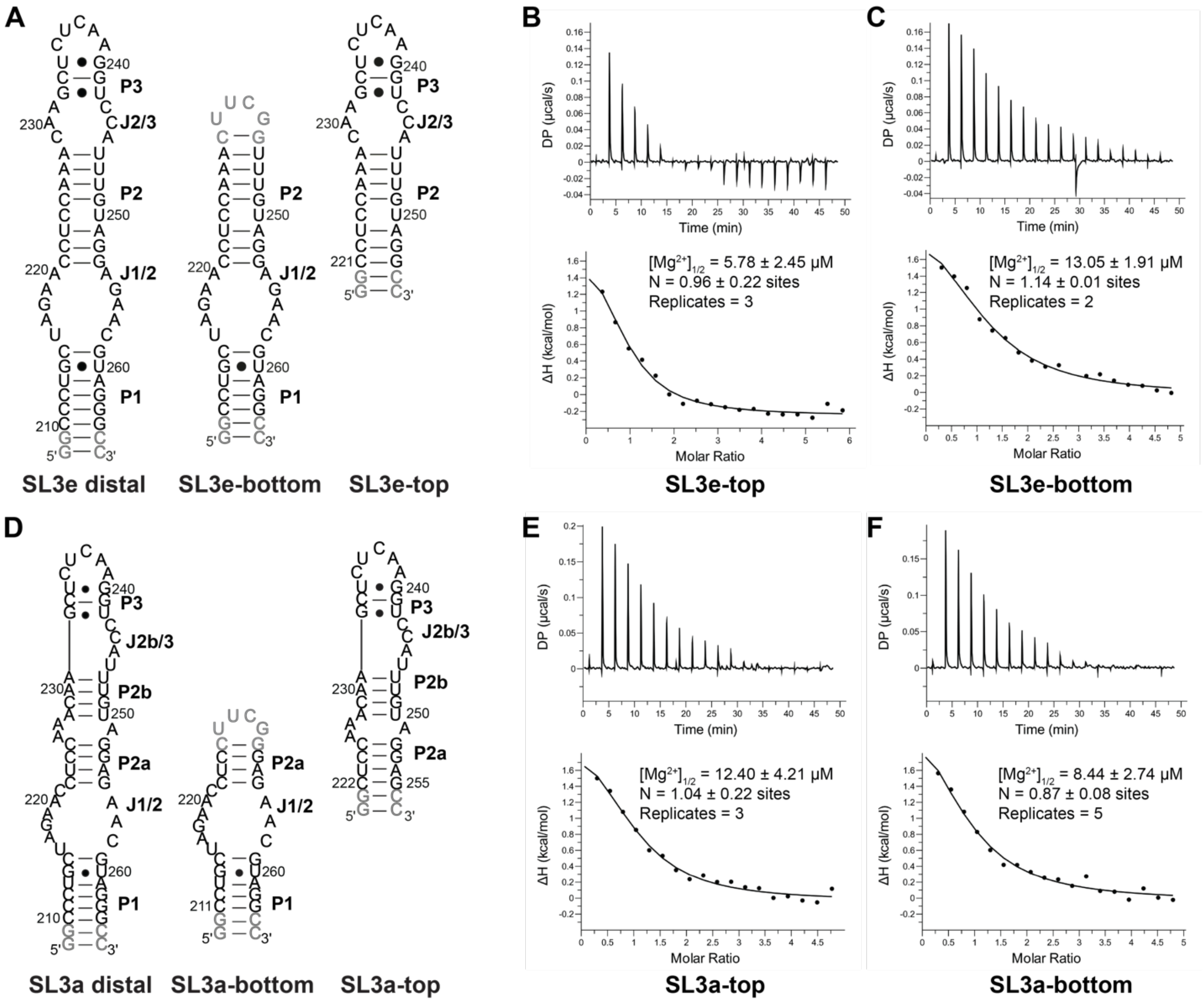


**Figure S4**: Representative $MgCl_2$ titration ITC thermograms of SL3 fragment constructs. A) Secondary structures of SL3 E-state and fragment constructs. Residues colored gray indicate non-native nucleotides introduced in the construct design, including tetraloop capping sequences and extra nucleotides at the 5′ and 3′ ends. B-C) Representative ITC thermograms for SL3e-top and SL3e-bottom respectively. D) Secondary structures of SL3 A-state and fragment constructs. E-F) Representative ITC thermograms for SL3a-top and SL3a-bottom respectively. The experiments were performed at 25 °C in a buffer of 100mM HEPES, pH8.0, and 60 mM NaOH for pH adjustment.

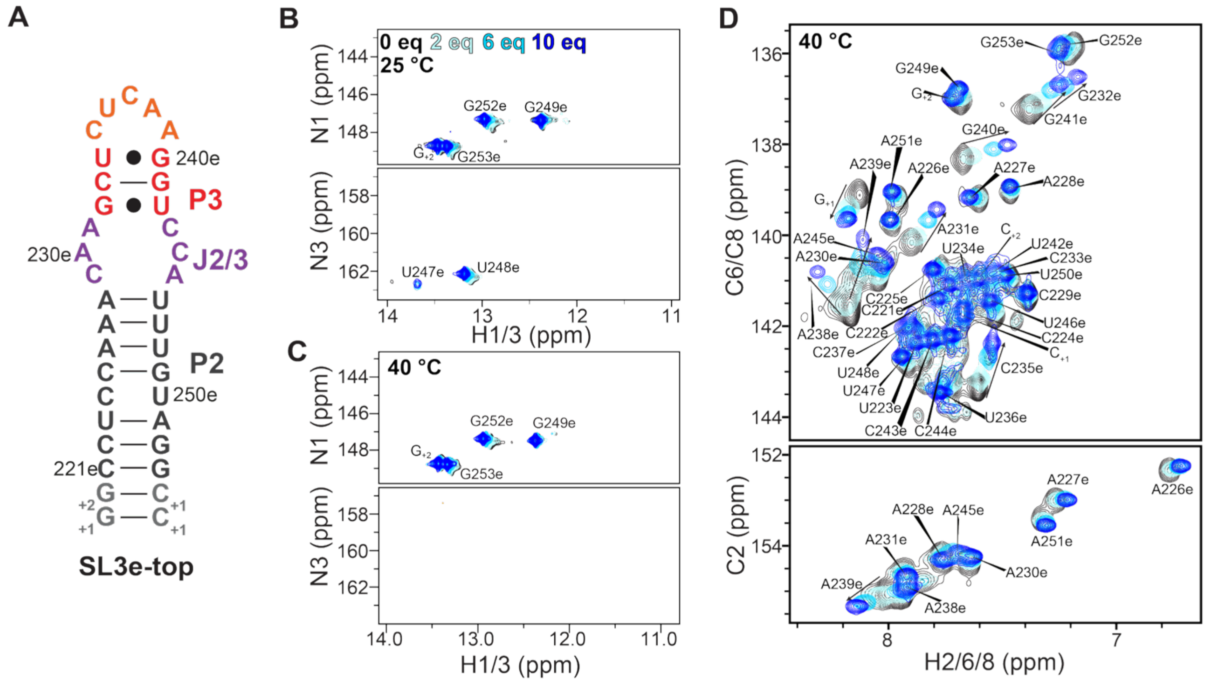


**Figure S5.** $^1$H-$^{15}$N and $^1$H-$^{13}$C HSQC spectra of SL3e-top $MgCl_2$ titration. A) Secondary structure of SL3e-top with residues in the stems and loops color-coded. Gray for P2 stem, purple for J2/3 loop, red for P3 stem, and coral for the apical loop. B) 2D $^1$H-$^{15}$N HSQC spectrum at 25 °C showing the disappearance of P3 stem and U223 resonances. C) 2D $^1$H-$^{15}$N HSQC spectrum at 40°C showing additional disappearance of U247 and U248 resonances. This behavior arises from increased solvent exchange of imino protons in RNA base pairs as temperature increases (1). D) 2D $^1$H-$^{13}$C HSQC spectrum at 40 °C showing chemical shift perturbations with increasing $Mg^{2+}$ concentration. Resonances in the absence of $MgCl_2$ are colored black, and with increasing $MgCl_2$ equivalents are colored varying shades of blue.

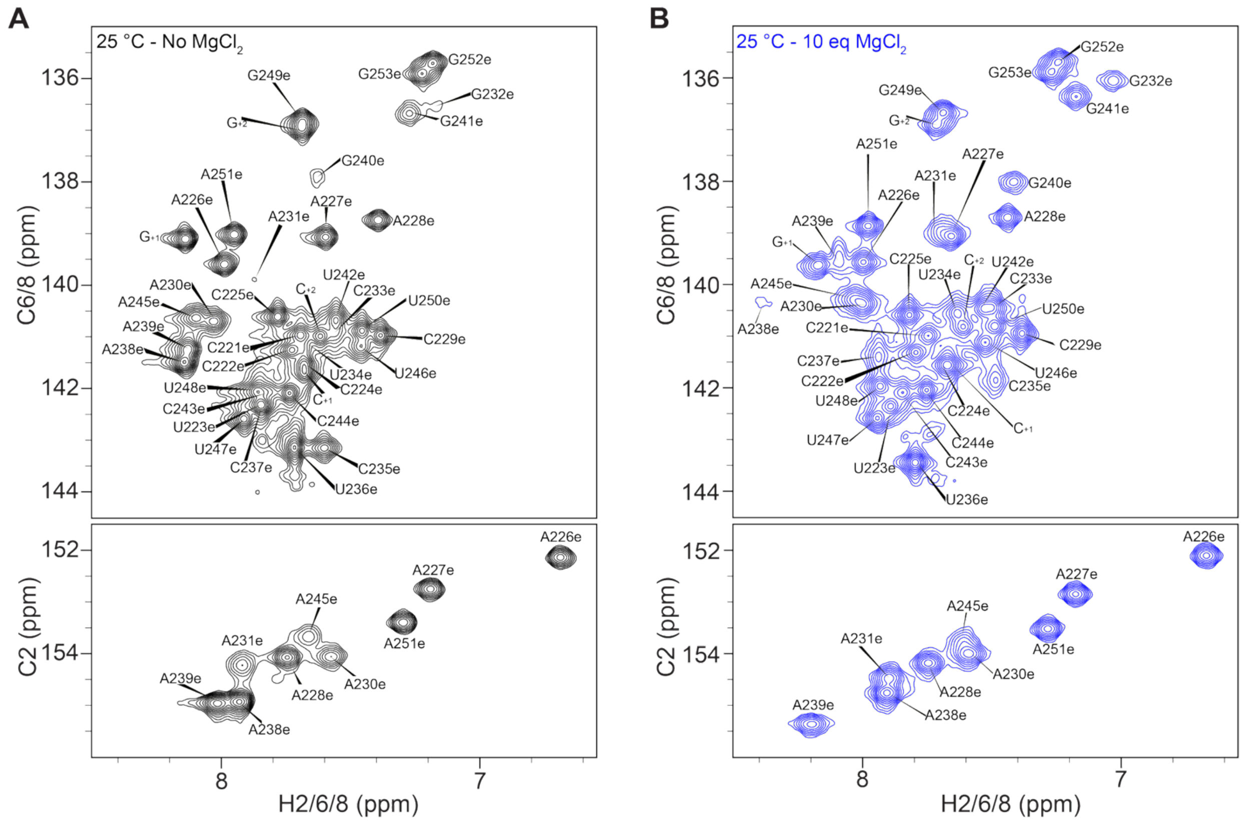


**Figure S6.** [1]H-[13]C HSQC spectra of SL3e-top at 25 °C. A) 2D [1]H-[13]C HSQC spectrum of SL3e-top in the absence of $MgCl_2$ colored black. B) 2D [1]H-[13]C HSQC spectrum of SL3e-top with 10 equivalents of $MgCl_2$ colored blue.

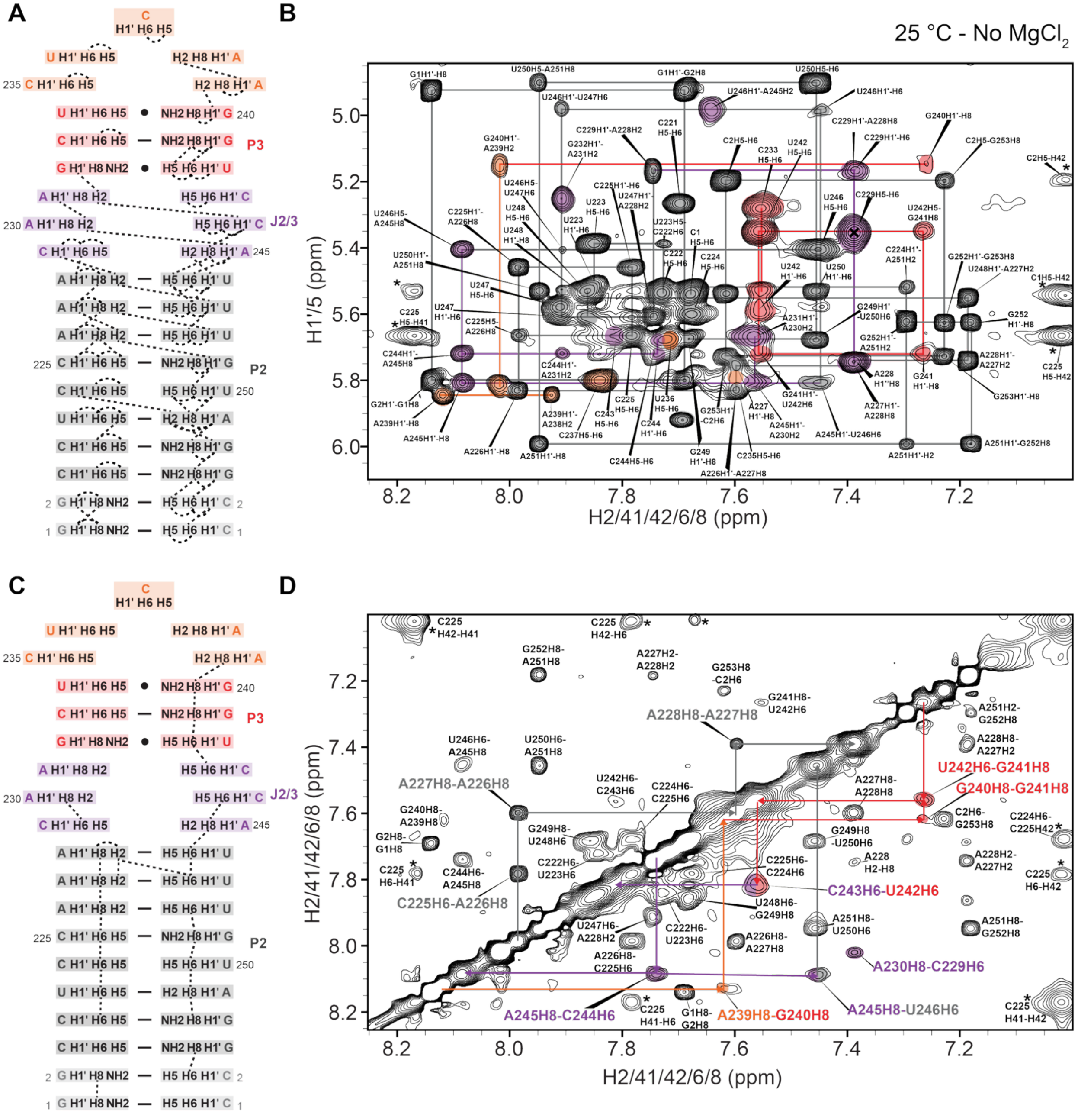


**Figure S7**. $^{1}$H-$^{1}$H NOESY spectra of SL3e-top in the absence of $MgCl_2$ at 25 ºC. A) Secondary structure of SL3e-top with dashed gray lines indicating sugar-to-base (H1'-H2/H6/H8) and base-to-base (H5-H6) connectivities observed in the absence of $MgCl_2$ as seen in the $^{1}$H-$^{1}$H NOESY spectrum shown in panel B. NOEs are colored according to secondary structure region, where P2 stem residues are colored gray, J2/3 loop residues are colored purple, P3 stem residues are colored red, and apical loop residues are colored orange. C) Secondary structure of SL3e-top with dashed gray lines indicating base-to-base (H6/H8-H6/H8) connectivities observed in the absence of $MgCl_2$. D) A region of $^{1}$H-$^{1}$H NOESY spectrum matching the connections in C. Dashed lines and labels are colored according to secondary structure region. Labels with increased font size highlight the base-to-base connections from A228-U246. With no $MgCl_2$, there are few connectivities in the J2/3 loop, P3 stem and the apical loop.

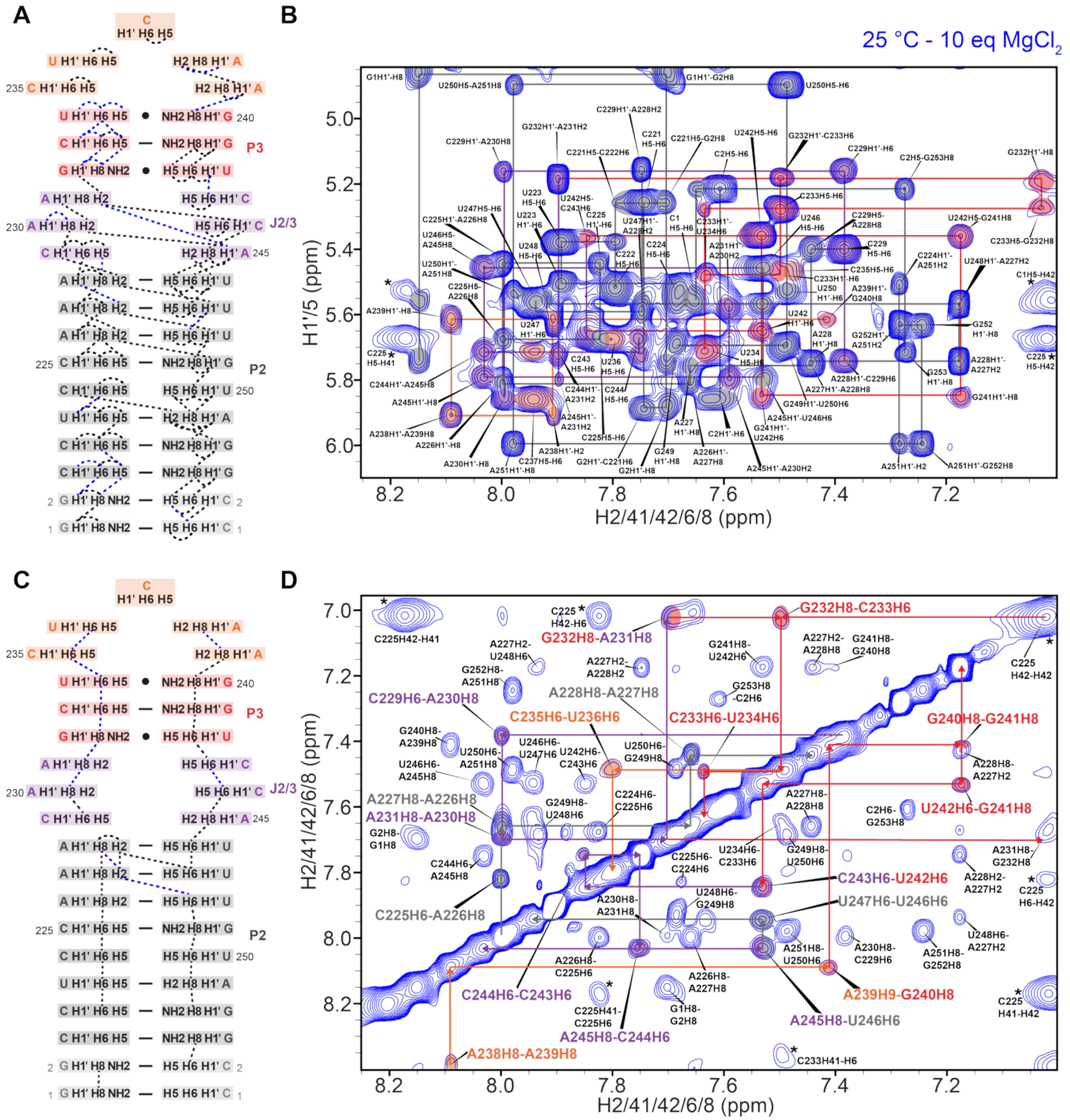


**Figure S8.** $^{1}H$-$^{1}H$ NOESY spectra of SL3e-top with 10 equivalents of $MgCl_2$ at 25 ºC. A) Secondary structure of SL3e-top with dashed lines indicating sugar-to-base (H1'-H2/H6/H8) and base-to-base (H5-H6) connectivities. Dashed gray lines indicate connectivities observed in the absence and presence of $MgCl_2$. Dashed blue lines indicate connectivities observed only in the presence of $MgCl_2$ observed in the $^{1}H$-$^{1}H$ NOESY spectrum in panel B. C) Secondary structure of SL3e-top indicating base-to-base (H6/H8-H6/H8) connections in the presence of 10 equivalents of $Mg^{2+}$ observed in the $^{1}H$-$^{1}H$ NOESY spectrum shown in panel D. Dashed lines and labels are colored according to secondary structure region. Labels with increased font size highlight the base-to-base connections from A228-U246. In the presence of 10 equivalents of $MgCl_2$, additional base-to-base connectivities are observed in the J2/3 loop, P3 stem, and apical loop.

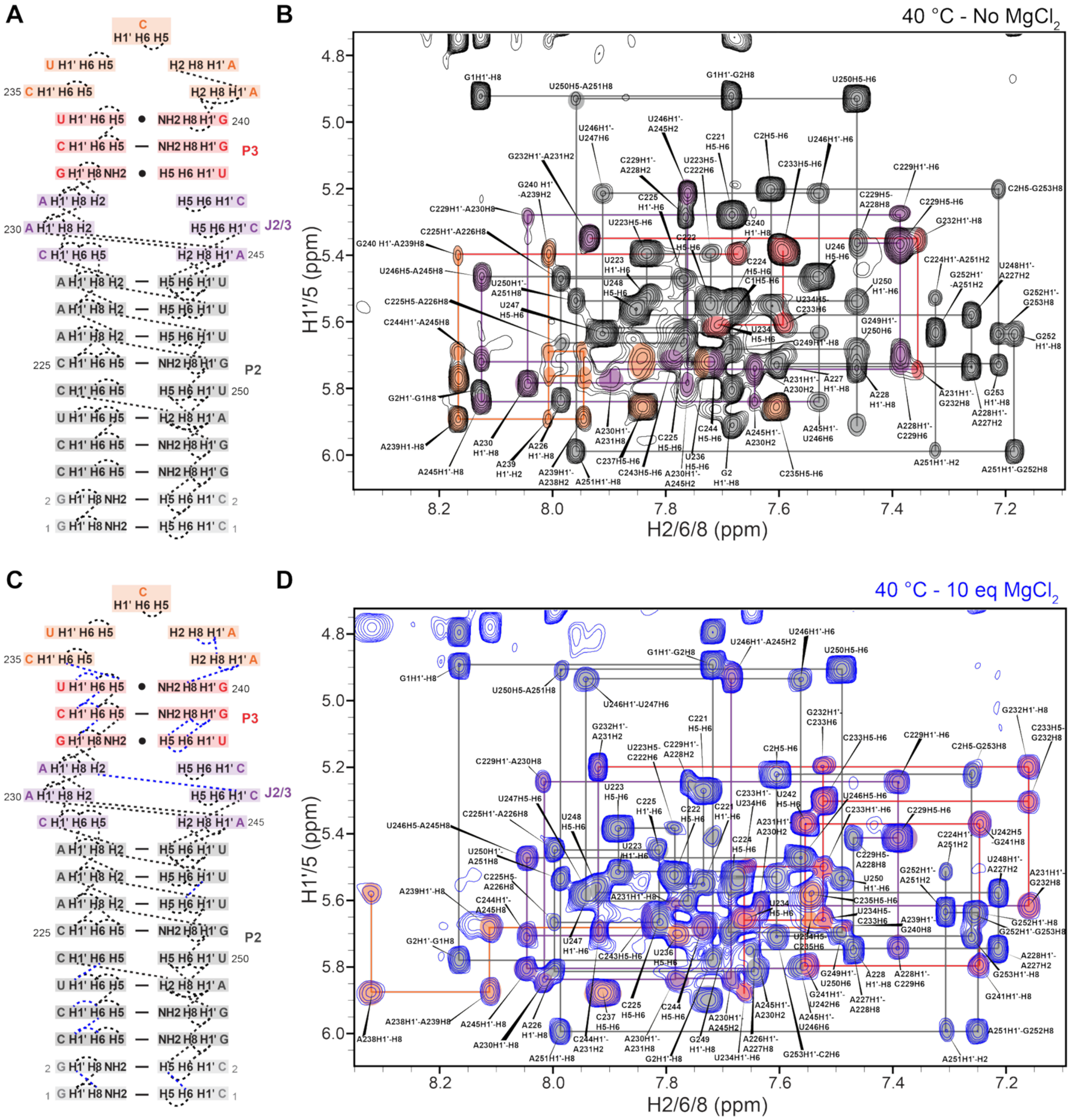


**Figure S9.** $^{1}H$-$^{1}H$ NOESY NMR spectra of SL3e-top in the absence and presence of $MgCl_2$ at 40 ºC. A) Secondary structure of SL3e-top with gray dashed lines indicating sugar-to-base (H1'-H2/H6/H8) and base-to-base (H5-H6) connectivities observed in the absence of $MgCl_2$. B) A region of $^{1}H$-$^{1}H$ NOESY spectrum showing the connections summarized in panel A. C) Secondary structure of SL3e-top with gray dashed lines indicating sugar-to-base (H1'-H2/H6/H8) and base-to-base (H5-H6) connectivities observed in both the absence and presence of $MgCl_2$. Dashed lines colored blue indicate connectivities observed only in the presence of $MgCl_2$. D) A region of $^{1}H$-$^{1}H$ NOESY spectrum matching the connections shown in C.

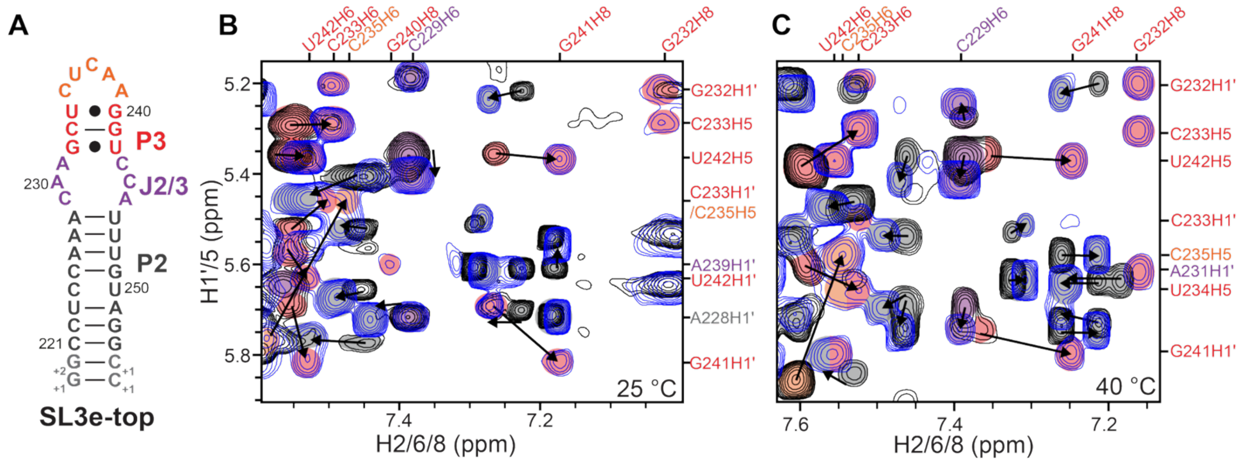


**Figure S10**. $^1H$-$^1H$ NOESY NMR spectral regions of SL3e-top supporting stabilized P3 stem with $Mg^{2+}$. A) Secondary structure of SL3e-top. B) A region of $^1H$-$^1H$ NOESY spectrum showing NOE connectivities in the P3 stem at 25 °C and C) 40 °C. NOEs observed in the absence of $MgCl_2$ are colored black, and NOEs observed in the presence of $MgCl_2$ are colored blue. The colored circles behind NOEs indicate the location of the residue on the secondary structure. In addition to chemical shift of NOEs generally changing with $Mg^{2+}$, there are appearance of new and well resolved NOEs for G232 and G241 at both B) 25 °C and C) 40 °C. This indicates stabilization of a helical geometry in the P3 stem.

**Supplemental references:**

1. Ferner, J., Villa, A., Duchardt, E., Widjajakusuma, E., Wohnert, J., Stock, G. and Schwalbe, H. (2008) NMR and MD studies of the temperature-dependent dynamics of RNA YNMG-tetraloops. *Nucleic Acids Res*, **36**, 1928-1940.
